# Functional and evolutionary impact of polymorphic inversions in the human genome

**DOI:** 10.1101/501981

**Authors:** Carla Giner-Delgado, Sergi Villatoro, Jon Lerga-Jaso, Magdalena Gayà-Vidal, Meritxell Oliva, David Castellano, Lorena Pantano, Bárbara D. Bitarello, David Izquierdo, Isaac Noguera, Iñigo Olalde, Alejandra Delprat, Antoine Blancher, Carles Lalueza-Fox, Tõnu Esko, Paul O’Reilly, Aida Andrés, Luca Ferretti, Marta Puig, Mario Cáceres

## Abstract

Inversions are one type of structural variants linked to phenotypic differences and adaptation in multiple organisms. However, there is still very little information about inversions in the human genome due to the difficulty of their detection. Here, thanks to the development of a new high-throughput genotyping method, we have performed a complete study of a representative set of 45 common human polymorphic inversions. Most inversions promoted by homologous recombination occur recurrently both in humans and great apes and, since they are not tagged by SNPs, they are missed by genome-wide association studies. Furthermore, there is an enrichment of inversions showing signatures of positive or balancing selection, diverse functional effects, such as gene disruption and gene-expression changes, or association with phenotypic traits. Therefore, our results indicate that the genome is more dynamic than previously thought and that human inversions have important functional and evolutionary consequences, making possible to determine for the first time their contribution to complex traits.

## INTRODUCTION

In the last decade a great effort has been devoted to characterizing the worldwide variation in the human genome (Altshuler et al., 2010; Francioli et al., 2014; Gurdasani et al., 2015; Sudmant et al., 2015; The 1000 Genomes Project Consortium, 2015; Walter et al., 2015). This information opens the door to determining the genetic basis of phenotypic traits and disease susceptibility. Nevertheless, despite the high initial expectations, a significant fraction of the genetic risk for common and complex diseases is still unexplained (Baker, 2012; Beckmann et al., 2007; Eichler et al., 2010; Manolio et al., 2009). Furthermore, not all types of variants have been studied at the same level of detail. In particular, inversions are a type of structural variant that changes the orientation of a genomic segment, usually without gain or loss of DNA, that have been mostly overlooked in humans (Alkan et al., 2011; Alves et al., 2012; Puig et al., 2015a).

The fact that they are balanced changes, together with the highly-identical inverted repeats (IRs) often found at their breakpoints, makes inversion detection very difficult, even with the newest sequencing methods based on short or long reads (Huddleston et al., 2017; Lucas-Lledó and Cáceres, 2013; Vicente-Salvador et al., 2017). Other techniques, like BioNano optical maps (Li et al., 2017) or Strand-seq (Sanders et al., 2016) promise to help in inversion discovery, but they are not suitable for high-throughput genotyping. Also, although inversion genotypes might be predicted based on SNP data (Cáceres and González, 2015; Cáceres et al., 2012; Ma and Amos, 2012; Salm et al., 2012), these methods can only detect certain types of inversions and the error rate can be high. Currently, a small number of inversions have been genotyped by FISH (Antonacci et al., 2009; Salm et al., 2012) and regular or inverse PCR (iPCR) (Aguado et al., 2014; Pang et al., 2013; Vicente-Salvador et al., 2017), which are labour intensive and are limited by the characteristics of the inversion IRs. Therefore, although inversions affect a significant fraction of the human genome (Martínez-Fundichely et al., 2014), only a few have been characterized in detail (Aguado et al., 2014; Antonacci et al., 2009; Pang et al., 2013; Puig et al., 2015b; Salm et al., 2012; Stefansson et al., 2005; Vicente-Salvador et al., 2017) and very little is known about the global frequency and distribution of inversions in human populations.

Actually, inversions have been a model in evolutionary biology for almost 90 years (Hoffmann and Rieseberg, 2008; Kirkpatrick, 2010) and there are numerous examples of their phenotypic consequences and adaptive significance in diverse organisms, from plants to birds (Jones et al., 2012; Joron et al., 2011; Küpper et al., 2016; Lowry and Willis, 2010; Thomas et al., 2008; Wellenreuther and Bernatchez, 2018). One of their main effects is related to recombination, since single crossovers within the inverted region in heterozygotes generate unbalanced gametes and at the same time the resulting inhibition of recombination could protect favorable allele combinations (Hoffmann and Rieseberg, 2008; Kirkpatrick, 2010). In addition, inversion breakpoints can alter directly the expression patterns of adjacent genes (Imsland et al., 2012; Puig et al., 2004, 2015b).

From the little information available, it is clear that inversions can have important functional consequences in humans (Puig et al., 2015a). Inversions are directly responsible of haemophilia A (Lakich et al., 1993) and they are associated with increased risk of neurodegenerative diseases (Baker et al., 1999; Myers et al., 2005; Pittman et al., 2006; Webb et al., 2008), autoimmune diseases (González et al., 2014; Namjou et al., 2014; Salm et al., 2012) or mental disorders (Collins et al., 2017; Okbay et al., 2016). The presence of an inversion could also predispose to other genomic rearrangements with negative phenotypic consequences in the offspring (Bayés et al., 2003; Gimelli et al., 2003; Hobart et al., 2010; Koolen et al., 2006). Moreover, there is evidence that a chr. 17 inversion increases the fertility of the carriers and has been positively selected in Europeans (Stefansson et al., 2005). Finally, inversions have been shown to affect gene expression (González et al., 2014; de Jong et al., 2012; Salm et al., 2012). However, most of these effects are associated with just two well-known inversions. Despite attempts to associate inversions with gene-expression and phenotypic variation in large datasets, the analyses have been limited exclusively to those with simple breakpoints, and only a couple of additional candidates have been identified so far (Chiang et al., 2017; Kehr et al., 2017; Sudmant et al., 2015). Thus, specific genotyping studies of a diverse range of inversions in a high number of individuals are necessary to have a more global idea of their functional and evolutionary impact in the human genome.

## RESULTS

### High-throughput genotyping of inversions

To overcome this limitation in the study of human inversions, here we have characterized in detail a representative set of 45 polymorphic paracentric inversions from the InvFEST database (Martínez-Fundichely et al., 2014), which corresponds approximately to half of the estimated number of real variants with >1% frequency in human populations (Sudmant et al., 2015) (Table S1, Figure S1). They are located throughout the genome (37 in the autosomes, seven in chr. X and one in chr. Y), with sizes ranging from 83 bp to 415 kb (median of 4.1 kb). In addition, 24 (53%) have been generated by non-allelic homologous recombination (NAHR) between >90% identical IRs (from 654 bp to 24.2 kb, with a median of 5.9 kb), whereas the rest (47%) were probably generated by non-homologous mechanisms (NH), including three with clean breakpoints and 18 with small deletions or insertions in the derived allele that may have been created in a single complex FoSTeS/MMBIR event (Sudmant et al., 2015; Vicente-Salvador et al., 2017). Three of those (HsInv0031, HsInv0045 and HsInv0098) have also short low-identity IRs (249-297 bp, 83.2-86.2% identity) in the ancestral orientation that are partially deleted in the derived orientation (Figure S1).

In order to genotype the inversions, we developed new high-throughput genotyping assays based on the multiplex ligation-dependent probe amplification (MLPA) technique (Schouten et al., 2002). For inversions with relatively simple breakpoint sequences (17), we carried out directly custom MLPA assays with minor modifications. However, for those with repetitive sequences at the breakpoints (24), which are very difficult to detect by most techniques, we devised and optimized a new method combining the principles of iPCR and MLPA named iMLPA. Briefly, we used two pairs of oligonucleotide probes for each inversion to interrogate the two alternative orientations, named as orientation 1 (*O1*) and orientation 2 (*O2*) (Figure 1). Four additional inversions not initially included in the MLPA-like assays were tested directly by multiplex PCR or iPCR (Table S1). The 45 inversions were genotyped in 551 individuals from seven worldwide populations used in HapMap and 1000 Genomes project (1000GP) (The 1000 Genomes Project Consortium, 2015; The International HapMap 3 Consortium, 2010) with an African (YRI, LWK), European (CEU, TSI), South-Asian (GIH) and East-Asian (CHB, JPT) ancestry, here referred as population groups (Table S2).

High-throughput inversion genotypes were then carefully validated through several analyses and quality controls (Figure 1), including comparison with 3,377 published genotypes (Table S1), identification of potential iPCR or iMLPA problems by analysis of restriction site polymorphisms, and analysis of the nucleotide variants associated to the inversion (see below). Genotypes with discrepancies or possible errors, plus other randomly selected genotypes, were tested by PCR or iPCR, including the whole set of 551 individuals for the three inversions that accumulated the highest error rate (HsInv0045, HsInv0055 and HsInv0340). All together, we checked 2,160 extra genotypes and generated a total of 24,355 inversion genotypes, with missing data (0.7%) accumulating mainly in specific problematic inversions or DNA samples (Figure S2A). The average error rate for the MLPA and iMLPA assays was also very low (0.1% and 0.9%, respectively), and again errors tended to accumulate in certain inversions due to restriction site polymorphisms or problems in MLPA probe amplification (Figure S2A), showing that the new inversion genotyping technique is very robust. Therefore, we have generated the largest and most accurate data set of different types of inversions in humans, including novel experimentally-resolved information on the worldwide distribution for all but three previously studied inversions (HsInv0030, HsInv0201 and HsInv0379).

In the final dataset, all inversions trios (30 CEU and 30 YRI) show correct genetic transmission and allele frequencies do not deviate from Hardy-Weinberg equilibrium in any population (*P* > 0.01). Minor allele frequencies (MAF) range globally from 0.5% to 49.7%, with 40 inversions (out of 45) present at >10% MAF in at least one population and most of them being spread in all population groups (Figure 2; Table S2). Only the four inversions with the lowest frequencies are polymorphic in a single population group (HsInv0097, HsInv0284 and HsInv0790 in Africa; HsInv0379 in East Asia), and eight more are fixed in one population group. On average, an African individual carries the *O2* allele for 28 inversions, whereas this number is slightly smaller for non-African individuals (24 inversions).

When compared to the only other available inversion population data set, only 14 (31%) of our 45 inversions were detected in the 1000GP (including 786 predicted inversions) (Sudmant et al., 2015), even though all of them were polymorphic in the studied populations. Moreover, considering the 434 individuals in common between both studies, a high error rate was observed in the 1000GP genotype data, with nine inversions with 2.5-71.5% incorrect genotypes (Figure S2B). Importantly, in the only two inversions with large IRs at the breakpoints identified by 1000GP (HsInv0241 and HsInv0374), the genotype agreement is extremely low (45.2% and 28.5%) and mostly due to a bias in favour of the reference genome orientation. This highlights the difficulty not only to detect inversions, but also to genotype them reliably from low-coverage short-read sequencing data, and emphasizes the importance of reliable genotypes for the characterization of human inversions.

**Figure 1.**
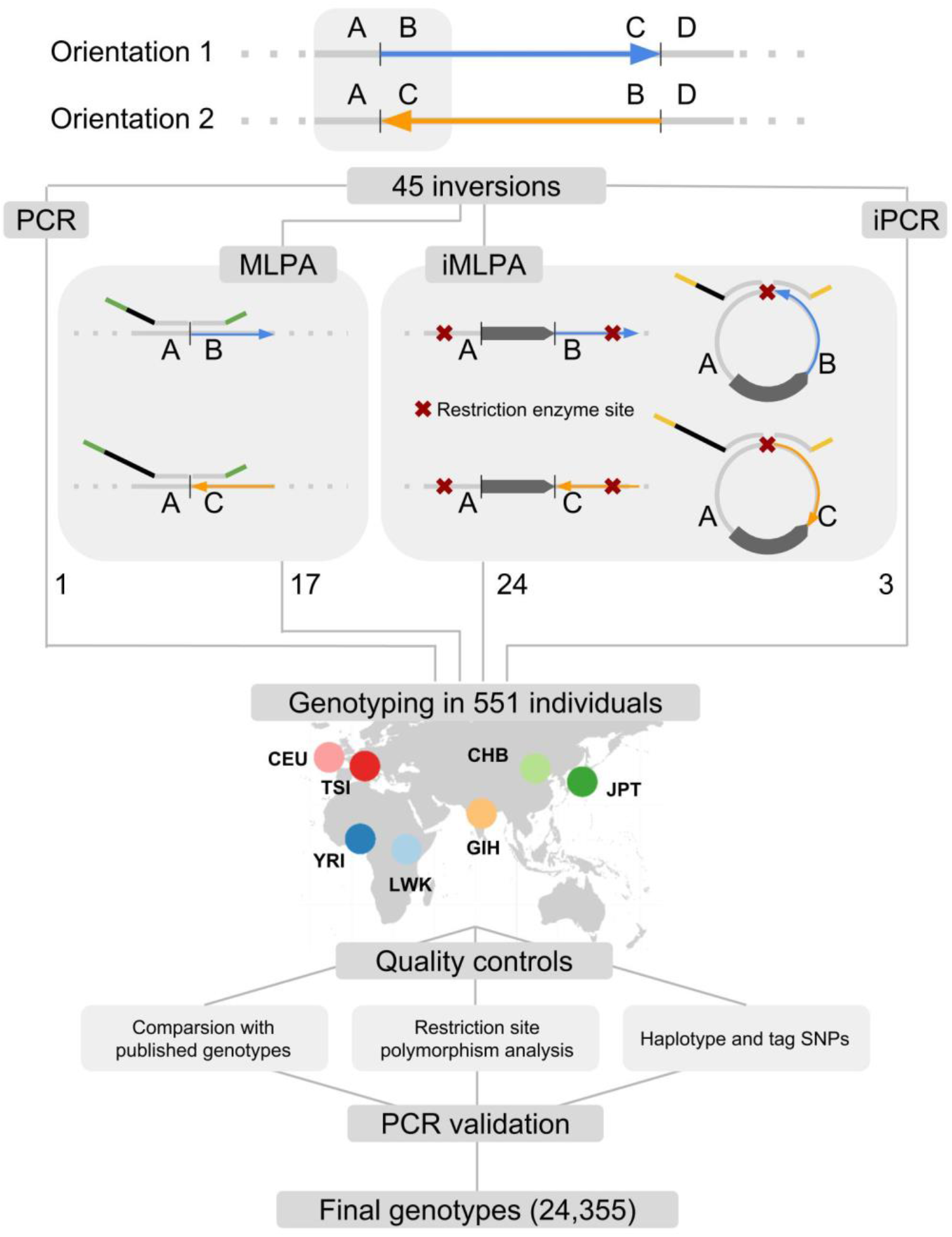
Schematic representation of inversion genotyping strategy. High-throughput genotyping of the 45 inversions in 551 individuals from different populations was done by MLPA (17). iMLPA (24), regular PCR (1) and iPCR (3). In MLPA and iMLPA the probes are represented in top of the genome sequence, including a sequence complementary to the genome (light grey), stuffer sequence (black) and common primers for amplification (green or yellow). Inverted repeats or other repetitive sequences at the breakpoints are represented as a dark pointed rectangle.

**Figure 2.**
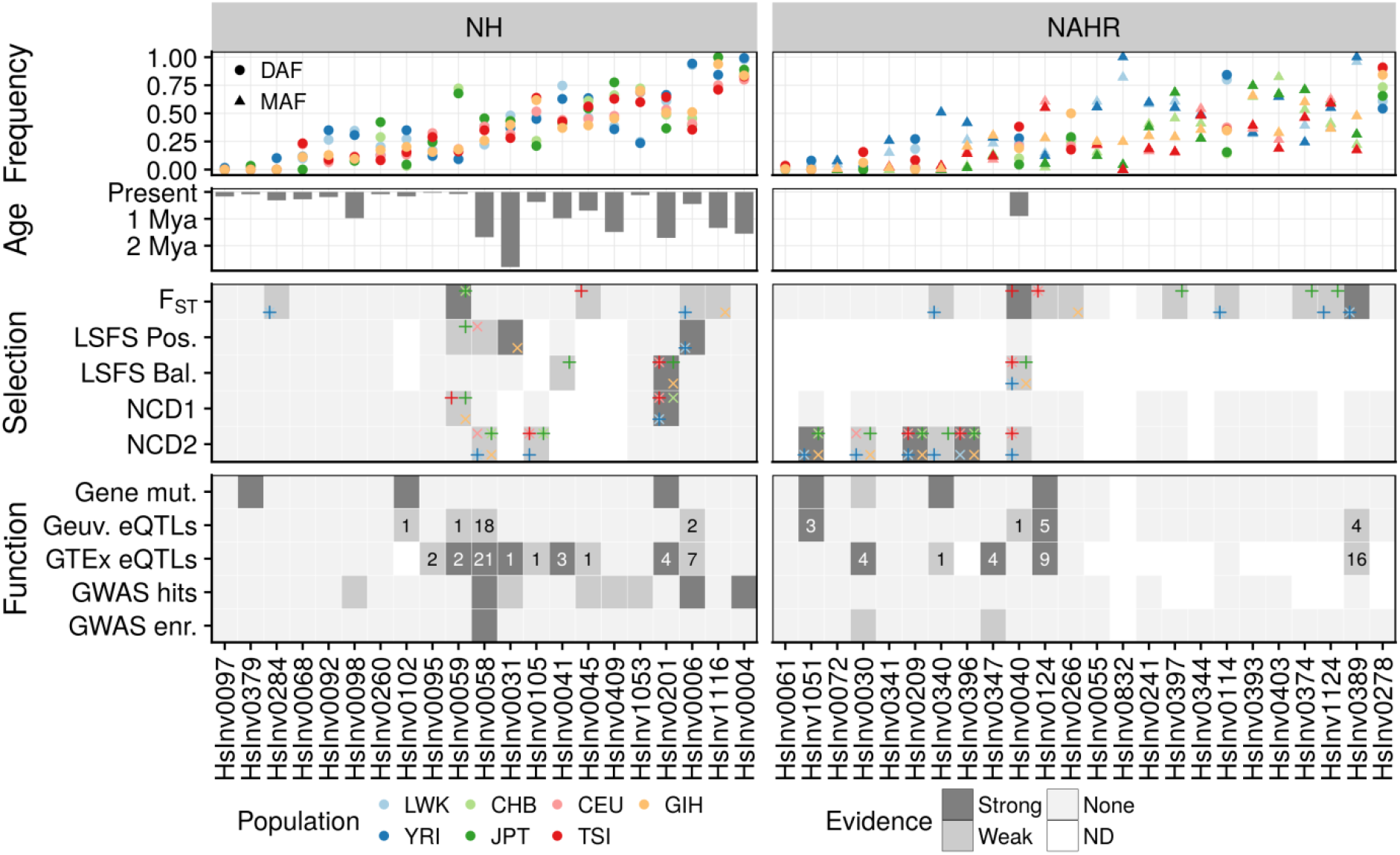
Evolutionary and functional information for human polymorphic inversions. First panel: inversion frequencies in the 480 unrelated individuals from seven populations, showing either the derived allele frequency (DAF) if the ancestral orientation is known or the minor allele frequency (MAF) according to the global frequency of the inversion otherwise (which enables the MAF to be higher than 0.5 in specific populations). Second panel: average inversion age obtained from divergence between orientations using three different substitution rate estimates. Third panel: summary of inversion selection signatures from FST, LSFS (positive or balancing selection) and NCD1 and NCD2 tests. Populations where the signal was detected are indicated in colors and criteria for classification of strong and weak selection evidences are explained in Methods. Fourth panel: functional effects of inversions summarizing direct gene mutation, which include gene disruption (strong) or exchange of genic sequences (weak), eQTLs in the GEUVADIS or GTEx datasets (showing the number of affected genes and labeled as strong if inversion is lead eQTL for at least one gene), and association with GWAS hits (strong if for one of the associations *P* < 1×10^−6^) or GWAS signal enrichment (strong if enrichment *P* < 0.01 in both GWAS databases).

### Association with nucleotide variants and haplotype distribution

Thanks to the accurate genotypes, we were able to explore the linkage disequilibrium (LD) between inversions and neighboring nucleotide variants (SNPs and small indels) from HapMap and 1000GP (The 1000 Genomes Project Consortium, 2015; The International HapMap 3 Consortium, 2010). We found consistent results in both datasets. While most NH inversions (20/21) have some variant in perfect LD (*r*^2^ = 1) either inside or up to 100 kb from the breakpoint, among the 24 NAHR inversions only HsInv0040 and HsInv1051 have at least one such variant (Figure 3A; Table S3). Maximum *r*^2^ values between the rest of NAHR inversions and other variants for the 434 individuals with 1000GP Phase 3 (Ph3) data ranged from 0.14 to 0.91, including only six NAHR inversions with *r*^2^ > 0.8 (Figure 3A).

Also, we checked the presence of shared polymorphic variants in both orientations based just in inversion and SNP (not including indels) genotypes. Most NH inversions do not have shared SNPs within the inverted region in accessible 1000GP positions or HapMap data, with the exception of a few individual SNPs that might be genotype errors or gene conversion events (Figure 3A; Table S3). Outside of the inversion, the proportion of shared SNPs increases progressively after the last fixed variant to a constant fraction of around 20% (Figure 3B). This makes it possible to define an extended area around the inversion of no or little recombination between chromosomes carrying different orientation (see Methods), which ranges from 0 to almost 20 kb on each side of the breakpoints with a median of ~3 kb (Table S3). In contrast, 20 of the 24 NAHR inversions have a considerable number of shared SNPs both throughout the inverted and flanking regions, with an average proportion of ~20% of SNPs being shared (Figure 3B). Since inversions inhibit recombination within the inverted region in heterozygotes and reduce the genetic flux between orientations (Hoffmann and Rieseberg, 2008; Kirkpatrick, 2010), this amount of shared variants was not expected and suggests the existence of other processes.

To better understand the relationship between the sequences of the two orientations, we combined nucleotide variants and inversion alleles in haplotypes, including both the inverted region and, when available, the region of recombination inhibition flanking the inversion. Next, we visualized the haplotype diversity and distribution across orientations and populations by generating haplotype networks and a new representation of haplotypes that integrates a hierarchical clustering and the differences between them (see Methods) (Figure S3). This analysis was focused on the 1000GP data, which provided higher resolution due to the larger number of variants, although consistent results were obtained with HapMap SNPs. After taking into account possible phasing errors, two clearly differentiated patterns could be observed that agree with the above results. In the 20 NH inversions with sequence variation information, the haplotypes of one of the orientations tend to cluster together, supporting a unique origin of the inversion (Figure S3). These haplotypes could be either clearly separated from those of the other orientation (*e.g*. HsInv1116, HsInv0004, HsInv0031, HsInv0058, HsInv0201 or HsInv0409), probably corresponding to old inversions that had time to diverge, or be clustered together with haplotypes carrying the other orientation, likely representing more recent or small inversions with few informative positions and little differentiation.

**Figure 3.**
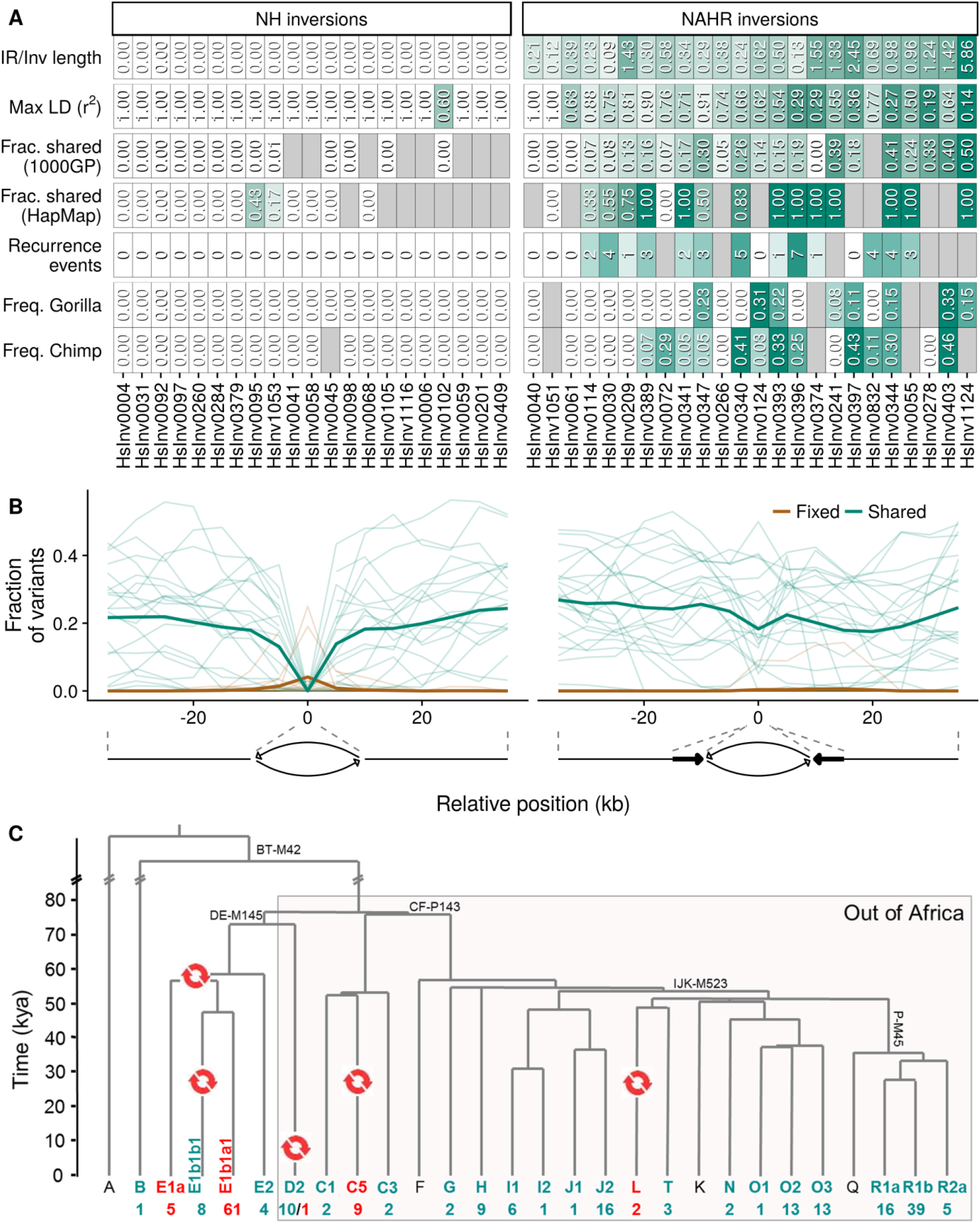
Evidence of unique or recurrent origin of inversions generated by non-homologous mechanisms (NH) and non-allelic homologous recombination (NAHR). **A**. Summary of inversion characteristics (ratio between breakpoint inverted repeat (IR) and inversion length), maximum linkage disequilibrium (LD) with neighboring variants in 1000GP and HapMap, fraction of shared SNPs inside the inverted region in 1000GP or HapMap, estimated number of recurrence events from haplotype analysis, and frequency of the derived (or *O2*) allele in non-human primates. Gray squares indicate values which could not be calculated. NAHR inversions show high heterogeneity in LD with nearby variants, proportion of shared polymorphisms between orientations, and frequency in other species, which suggest different levels of recurrence. **B**. Distribution of fixed (brown) and shared (green) variants between the two orientations estimated from inversion genotypes and 1000GP data. The shared/fixed fraction with respect to the total number of polymorphic variants was computed for the whole inverted region plus 10-kb overlapping windows with a 5-kb step size in the flanking regions, and the horizontal axis represents the distance of the window central position to the inversion breakpoints (indicated by dashed lines). Thin lines represent individual inversions and thick ones represent the average for each inversion class. **C**. Overview of chr. Y phylogenetic tree in humans showing five different possible inversion events in the HsInv0832 region (red arrows). Divergence dates (left scale) and tree topology are based on Poznik *al*. (2016) (Poznik et al., 2016). Chr. Y haplogroups and the number of males genotyped for HsInv0832 are indicated at the bottom of each branch, with green and red colors representing the *O1* or *O2* orientation, respectively. Only the main branches and those including genotyped individuals are shown, with some characteristic mutations indicated in the tree. For haplogroup E, one of the two possible scenarios is represented, with the alternative being two inversion events both in E1a and E1B1a1 haplogroups.

In contrast, this is true for only three of the NAHR inversions (HsInv0040, HsInv0061 and HsInv1051), with the other 21 having *O1* and *O2* haplotypes mixed throughout the network and hierarchical cluster, including in many cases identical haplotypes with both orientations (Figure S3). Such pattern is consistent with a multiple origin of these inversions and explains the absence of fixed SNPs and the high number of shared SNPs between orientations. Based on the results of the different analyses, we inferred a conservative estimate of the minimum number of recurrent inversion events and approximately map their distribution in human populations. However, this relies on the existence of differentiated haplotype clusters in which the existence of the other orientation cannot be easily explained by other factors (such as gene conversion or genotype/phasing errors of a few variants). Therefore, it is not possible to detect more than one event within a cluster and there is a bias to predict more potential recurrence in larger inversions with more variants, so it is necessary to interpret the results with caution. In particular, in six of the smallest NAHR inversions, *O1* and *O2* haplotypes are too similar to identify individual recurrence events (Table S4). In two others (HsInv0124, HsInv0397), most *O1* and *O2* haplotypes belong to the same big cluster with just few differences between them and no clear recurrence can be identified.

Another problem is that haplotype recombination generates mixed sequences between two groups and makes it difficult to accurately quantify recurrence. Thus, a nice example is inversion HsInv0832 in the haploid chr. Y, in which there is no recombination or need to infer haplotypes by phasing. Since the chr. Y genealogical tree is already known, we used available haplogroup information of 232 males (Table S5) to identify five independent inversion events in the last ~60,000 years, including two in African haplogoup E and three other inversions in Asian haplogoups C, D, and L (Figure 3C). This results in an inversion rate of 5.34 × 10^−5^ per generation (see Methods), which is ~1,000 times higher than that of single bases.

Overall, for the inversions in which it is possible to quantify recurrence, we estimated a total of 40 additional inversion and re-inversion events (ranging for each inversion from 1 to 7 with an average of 3), (Table S5). Of those, 12 are distributed globally and could have occurred before the out-of-Africa migration, 17 are restricted to African individuals, and 11 have probably appeared more recently in non-African populations. The fact that many of the recurrence events are shared by several individuals indicates that they are real evolutionary events and not artefacts from LCL cell culture. In addition, as part of the validation of inversion genotypes, all the recurrence events were confirmed by checking at least one of the supporting individuals by PCR/iPCR, plus many additional individuals of other inversions having a high proportion of shared SNPs. Thus, our results extend considerably the previous analysis in just the CEU population (Aguado et al., 2014; Vicente-Salvador et al., 2017), with nine more inversion events detected in the five inversions originally predicted as recurrent in humans (HsInv0030, HsInv0341, HsInv0344, HsInv0389 and HsInv0393) and 7 of the 12 inversions considered to be unique or lacking information in CEU being now recurrent as well.

### Ancestral orientation and inversion age

To get a better idea of the mutations directionality, we also checked the ancestral orientation of the inversion regions. This had been described for 32 of the analyzed inversions (Aguado et al., 2014; Puig et al., 2015b; Vicente-Salvador et al., 2017). However, we complemented and expanded the published data by experimentally genotyping 42 inversions in a panel of 23 chimpanzees (40 inversions) and 7 gorillas (41 inversions) with either the human or modified assays (Table S6). The orientation of the inverted region was also assessed in available genome assemblies of four nonhuman primate species, including orangutan and rhesus macaque (Table S7). In total, we could infer the ancestral allele for 29 inversions, with half showing the ancestral (15) or the derived (14) allele in the reference human genome. Of those, the 21 NH inversions had consistent orientations in all the genotyped samples, the non-human genomes analyzed, and the published data. Moreover, this was also consistent with the proposed mechanism of generation, with additional deletions and insertions occurring in the derived allele for inversions mediated by more complex events (Table S1).

In contrast, 14 of the 22 NAHR inversions experimentally genotyped were polymorphic in at least one of the ape species, with opposite orientations in different primate assemblies for several of them. Thus, the ancestral orientation could not be determined (Table S7). Of those, six inversions were polymorphic in both species, six only in chimpanzees and two only in gorillas, and in all of them the genotypes were confirmed by PCRs of at least one breakpoint. This agrees with the previous results in a much smaller set of samples (Aguado et al., 2014; Vicente-Salvador et al., 2017), but we found five additional polymorphic inversions in each species. In fact, the lower number of polymorphic inversions in gorilla could be due to the reduced sample, which suggests that as more individuals are analyzed, more of these inversion regions might be identified as polymorphic in non-human primates.

In terms of the allele frequency for the 12 polymorphic inversions in chimpanzees, the correlation with that in humans is quite low (*r*^2^ = 0.055), and in seven inversions the minor allele is different in both species. Given the three species divergence times, the most likely scenario is that shared inversions have appeared independently in the chimpanzee and gorilla lineages, providing additional support for the evidence of recurrence from human data (Figure 3A).

Moreover, we took advantage of available ancient Hominin genomes to check the presence of the studied inversions. Specifically, we looked for the breakpoint sequences in both orientations of 19 NH inversions without IRs in Neanderthal, Denisovan and two modern human individuals of 5,300 (Otzi) and 7,000 (La Braña 1) years (Table S7). Five inversions (HsInv0004, HsInv0006, HsInv0201, HsInv0409 and HsInv1116) showed the derived orientation in the Neanderthal or Denisovan genomes (including two with the derived orientation in both), which suggests that they appeared before the divergence of the most recent *Homo* groups, around 550,000-750,000 years ago (ya) (Prüfer et al., 2014). These inversions are distributed in African and non-African populations and it is very unlikely that they are the result of introgression. In the two early modern human genomes analyzed, we confirmed the existence of the derived allele in eight inversions, all corresponding to those with a global frequency over 0.15.

Finally, we dated more precisely 22 inversions with a unique origin based on the divergence between orientations from 1000GP data using three different substitution rate estimates (Figure 2; Table S7). As expected from the ancient genomes results and the haplotype analysis, six inversions were estimated to have appeared more than 1 Mya, including four with the derived orientation in Neanderthal or Denisovan, plus HsInv0031 and HsInv0058. For HsInv0006, the estimated age (407,795-495,470 ya) is slightly more recent than the Neanderthal-Denisova and modern humans divergence, but it is very close to its lower bound. Ages of the rest of inversions are consistent with their geographical distribution, with inversions restricted to either African (HsInv0097 and HsInv0284) or non-African populations (HsInv0379) having a relatively recent origin. Only in two cosmopolitan inversions age estimates are lower than population split times (HsInv1053 with a negative age estimate and 22,582-41,258 ya for HsInv0095), probably due to an underestimate of the divergence between orientations caused by the limited sequence information available and the presence of shared SNPs.

### Analysis of inversion frequencies

To evaluate whether there are selective pressures acting globally on inversions, we first compared the inversion frequency spectrum with that of SNPs sampled according to neutral expectations (see Methods). While the ascertainment bias associated with inversion detection predicts enrichment of high-frequency inversions (~0.20 expected MAF), it does not explain the observed frequencies, which tend to be higher than expected in all populations (Figure 4A). Stratifying inversions by the mechanism of generation and using either MAF or DAF (if ancestral orientation could be determined), demonstrated that the increased frequency is evident only in NAHR inversions (Figure 4A). Potential explanations for this are that the detection bias associated to IRs has been underestimated or that there are other mutational or selective forces affecting these inversions.

**Figure 4.**
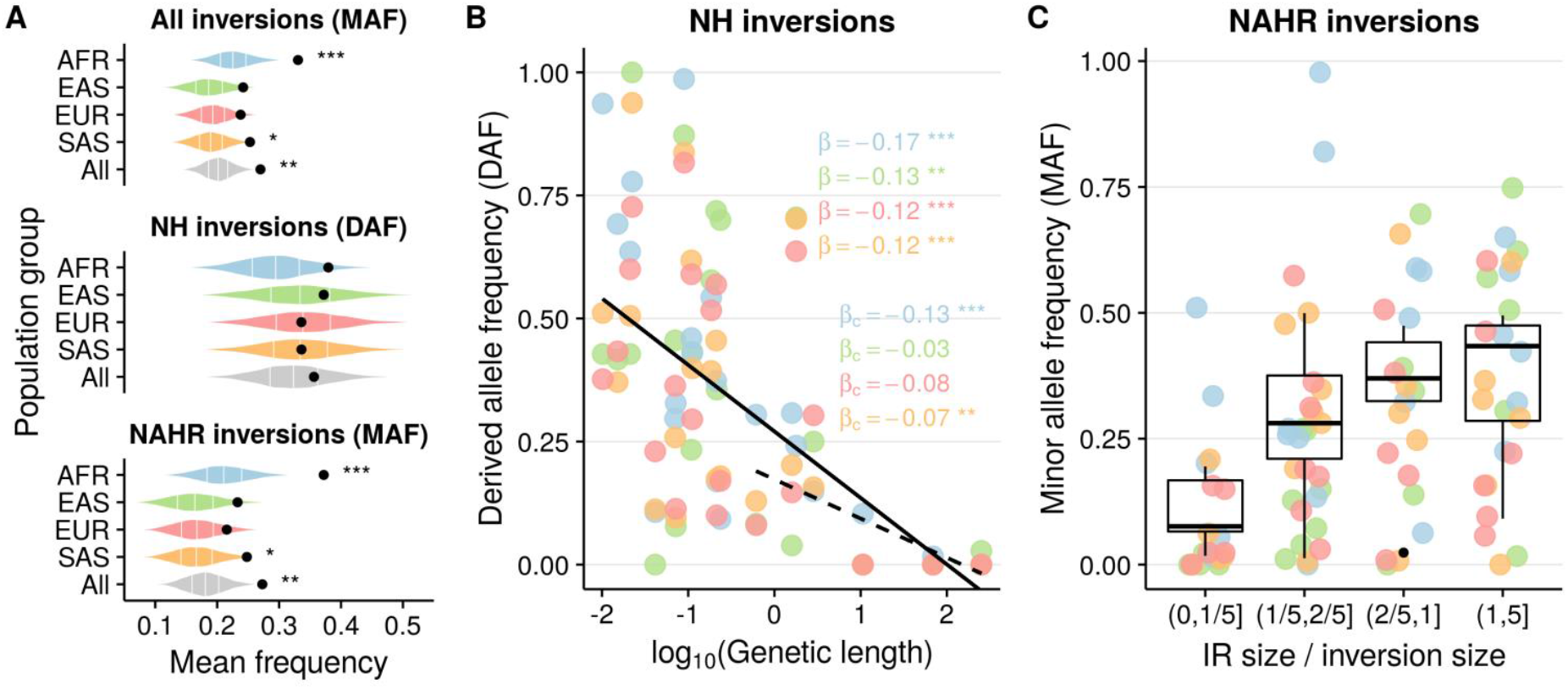
Determinants of inversion frequency in human populations. **A**. Observed mean inversion frequency per population group and mechanism of generation (using DAF only when known for all inversions in the group and MAF otherwise) compared with that expected from the detection method simulations using SNPs (violin plots with marked quartiles). Mean frequencies were compared with a two-sided permutation test. **B**. Inverse logarithmic robust regression between DAF of NH inversions and genetic length (measured in 4N_e_r units), showing a significant trend for all inversions (solid line; ß regression coefficient) and for inversions larger than 2 kb to correct in part the detection bias (dashed line; ß_c_ regression coefficient). **C**. Boxplots showing a relationship between the frequency of the minor allele in all populations together and IR/inversion length ratio for NAHR inversions. In **B** and **C** dots indicate the frequency of each inversion per population group, represented with the same colors as in **A**. *, *P* < 0.05; **, *P* < 0.01; and ***, *P* < 0.001.

Next, we investigated the impact of diverse genomic variables on autosomal and chr. X inversion frequencies. The most significant predictor is the inversion genetic length, which is negatively correlated with DAF of NH inversions, even after controlling for the detection bias, and explains 23-55% of the frequency variance in the different population groups (Figure 4B), followed by physical length. This relationship is much weaker for NAHR inversions, with 5-14% of MAF variance explained by inversion genetic length. In fact, physical length of NAHR inversions increases with IR length (*R*^2^ = 0.24, *P* = 0.0076), and the ratio between IRs and inversion length appeared as the main factor, explaining 13-45% of MAF variance (Figure 4C). Altogether, these results suggest that there are two mechanisms driving inversion frequency in different directions: (1) the potential negative selection against long inversions resulting from the generation of unbalanced gametes by recombination (Hoffmann and Rieseberg, 2008; Kirkpatrick, 2010); and (2) a recurrent back-and-forth generation process for inversions with high IR-inversion size ratio, converging to equilibrium frequencies of ~0.5.

### Selection on human inversions

We also used several complementary strategies to investigate signals of natural selection acting on specific inversions. We considered two different selective scenarios: (1) short-term positive selection leading to a rapid increase in allele frequency, possibly in one or a few populations; and (2) long-term balancing selection maintaining a polymorphism in several populations.

First, we measured inversion frequency differences between populations using the fixation index (F_ST_), which can identify positive selection in specific regions. The global F_ST_ value was 0.11 for autosomal inversions, 0.21 for chr. X inversions, and 0.73 for the one in chr. Y, with the largest frequency variation between continents, as expected. In particular, we found three inversions within the top 1% of the F_ST_ distributions derived from SNPs with the same frequency (Figure 2; Table S8), including large frequency differences between European populations (HsInv0040) or population groups (with high frequency in Africa for HsInv0389 and in East Asia for HsInv0059). Eleven more inversions fall within the top 5% of the empirical F_ST_ distribution, and were considered to have weak evidence of selection (Figure 2; Table S8).

Second, we applied a novel neutrality test based on the frequency spectrum of linked sites (LSFS), which is well suited to detect deviations from neutrality in low-recombination regions, such as inversions (see Methods). We used optimised tests to identify positive and balancing selection and significance was assessed empirically using the genome-wide distribution from autosomes (see Methods). Since phased sequences are required, only 18 autosomal inversions with perfect tag variants near the inverted region were analyzed, including most NH inversions and HsInv0040 (Figure 2; Table S8). The strongest signals (*P* < 0.01) were in HsInv0201 for balancing selection and inHsInv0006 and HsInv0031 for positive selection. In addition, there are four other inversions with weaker evidence of balancing or positive selection. Consistent with the F_ST_ results, in HsInv0006 and HsInv0059 LSFS positive selection signals are detected in those populations with increased DAF (Figure 2; Table S8).

Finally, as independent confirmation of balancing selection signatures, we used the recently developed non-central deviation statistics, NCD1 and NCD2 (Bitarello et al., 2018). NCD1 detects site frequency spectrum shifts towards an equilibrium frequency as expected under balancing selection, whereas NCD2 incorporates also information on polymorphism density and is most powerful to detect long-term balancing selection (Bitarello et al., 2018). However, the results of these tests summarize the data of all the SNPs in a region and are not necessarily linked to the inversion, as in the previous cases. An empirical *P*-value was assigned for each inversion region and population by comparing them with a genome-wide distribution (Table S8). Focusing on signals detected in at least three populations, we found respectively four and six inversions with strong and weak signatures of balancing selection for NCD1 or NCD2 (Figure 2). Many of these candidates could not be analyzed with the LSFS method because of the lack of tag SNPs, but consistent results were found for HsInv0201. Other cases correspond to low frequency inversions, such as HsInv1051 and HsInv0209, and potential selection seems more related to the region than the inversion itself. Another example is HsInv0058, in which the signal is likely due to the different haplotypes of the MHC region (see below).

### Effect of inversions on genes and gene expression

Inversions can have important effects on genes, as previously described for some of those analyzed here (Aguado et al., 2014; Pang et al., 2013; Puig et al., 2015a, 2015b; Sudmant et al., 2015; Vicente-Salvador et al., 2017). Almost half of our inversions (21/45) are located in intergenic regions, although seven of them are within 20 kb of protein-coding genes or long non-coding transcripts (Table S9). In addition, three inversions invert genes and eight are located within introns. In seven other inversions, genes overlap the two highly-identical IRs at the breakpoints and there might be an exchange of gene sequences, as happens in HsInv0030 (Pang et al., 2013; Vicente-Salvador et al., 2017) and probably also in HsInv0241, HsInv0396 and HsInv1124, since the genes span most of the IRs. Finally, there are six inversions that affect genes more directly (Figure 2), involving the inversion or deletion of an internal exon (HsInv0102, HsInv0201) and the disruption of an alternative isoform (HsInv0124) or the whole gene (HsInv0340, HsInv0379, HsInv1051), which in the case of HsInv0379 resulted in the loss of its expression and the generation of a new fusion transcript (Puig et al., 2015b).

We measured the effect of the 42 autosomal and chr. X inversions with MAF > 0.01 on expression of nearby genes using lymphoblastoid cell line (LCL) transcriptome data of 445 individuals from European (358) and African (87) origin of the GEUVADIS consortium (Lappalainen et al., 2013). To increase statistical power and reliability, the analysis was replicated in two data sets: (1) 173 CEU, TSI and YRI individuals with inversion genotypes; and (2) an extended set with 272 additional European and YRI individuals in which the genotypes of 33 inversions could be imputed through correlated neighboring variants (Figure S4A). cis-eQTL associations were performed by linear regression between inversion genotypes and expression profiles, testing a total of 850 genes and 3,318 transcripts with an inversion within 1 Mb of the transcription start site (TSS). Considering the largest sample size for each inversion, we uncovered eight inversions significantly associated with LCL expression of 27 genes and 44 transcripts (FDR 5%) (Figure S4B; Table S10). For inversions analyzed in the experimentally-genotyped and imputed data sets the results were highly concordant, since virtually all gene and transcript inversion eQTLs were replicated (7/7 and 11/12, respectively) and effects were always in the same direction (Figure S4C). As negative control, a null distribution of no association was observed by permuting inversion genotypes relative to expression levels (Figure S4D). Moreover, significant expression effects were robust when applying different analysis approaches (see Figure S4E-F and Methods). Interestingly, inversions acting as eQTLs tended to locate significantly closer to the TSSs of the differentially expressed genes (<100 kb) in comparison to all tests performed (Figure S4D). Therefore, these analyses strongly support the inversion-expression associations detected in LCLs.

Next, we examined the effect of inversions on gene expression in other tissues through variants in LD (*r*^2^ ≥ 0.8) already reported as eQTLs in the GTEx project (GTEx Consortium, 2013, 2017). We found 62 genes with eQTLs in different tissues highly linked to 11 of the 26 analyzed inversions, including seven for which no expression differences were found in LCL data (Table S11). Good consistency was also found with the previous results, with 17/27 of inversion-gene associations in LCLs that could be checked being also identified in GTEx data (Figure S5). Although this analysis provides an initial estimation of inversions contribution to expression variation, most of the recurrent inversions could not be analyzed in other tissues. By searching for eQTL signals in moderate LD (*r*^2^ ≥ 0.6) with those inversions, we found evidence of additional potential expression effects for HsInv0124 and novel expression changes in the genes affected by two inversions (Figure S5): HsInv0030, which exchanges the first exon and promoter of chymotrypsinogen precursor genes *CTRB1* and *CTRB2* only expressed in pancreas (Pang et al., 2013; Vicente-Salvador et al., 2017), and HsInv0340, which disrupts the long non-coding gene *LINC00395* expressed in testis.

To assess whether inversions or other variants in high LD were the main cause of the observed expression changes, we performed a joint eQTL analysis in LCLs including our inversions together with SNPs, indels and structural variants from the 1000GP (Sudmant et al., 2015; The 1000 Genomes Project Consortium, 2015). We defined the lead marker as the most likely causal locus for each eQTL discovered, resulting in inversions HsInv0124 and HsInv1051 showing the highest association signal for two genes and three transcripts (Figure 5A). We also checked the association of inversions with neighboring SNPs and the top lead eQTLs detected by the GTEx project. Six inversions show the highest LD (*r*^2^ ≥ 0.9) with variants reported as first or second lead eQTL in a given tissue (Figure S6). In general, some of the strongest effects are related to inversions breaking or affecting exonic sequences, although the consequences can be complex and need to be investigated in detail. For example, in HsInv1051, which breaks the *CCDC144B* gene, the expression increase of specific alternatively-spliced isoforms (Figure 5A) is really due to the creation of a fusion transcript with new sequences at 3’ that loses two thirds of the gene exons (Figure 5B; Figure S7). For recurrent inversions HsInv0124 and HsInv0030, the increase in the eQTL-gene association with LD with the inversions indicates that they could be causing the expression changes (Figure 5C). In fact, HsInv0124 is the lead variant for the antisense RNAs *RP11-326C3.7* and *RP11-326C3.11* at gene and transcript level, which overlap respectively the *IFITM2* and *IFITM3* genes located at the breakpoints, and it has opposite effects in the two pairs of overlapping genes (Table S10; Table S11). Also, HsInv0102 clearly affects those *RHOH* isoforms that express the alternative non-coding exon inverted by the inversion, but its effect is partially masked by a more frequent lead eQTL (SNP rs7699141) that acts in the same direction. On the other hand, HsInv0058 is associated to chr. 6 MHC haplotypes APD, COX, DBB, QBL and SSTO, which extend ~4 Mb and are known to harbor important functional differences (Horton et al., 2008). In this case, the lower linkage of the inversion with lead eQTLs suggests that other variants in these haplotypes are really responsible of the observed effects.

**Figure 5.**
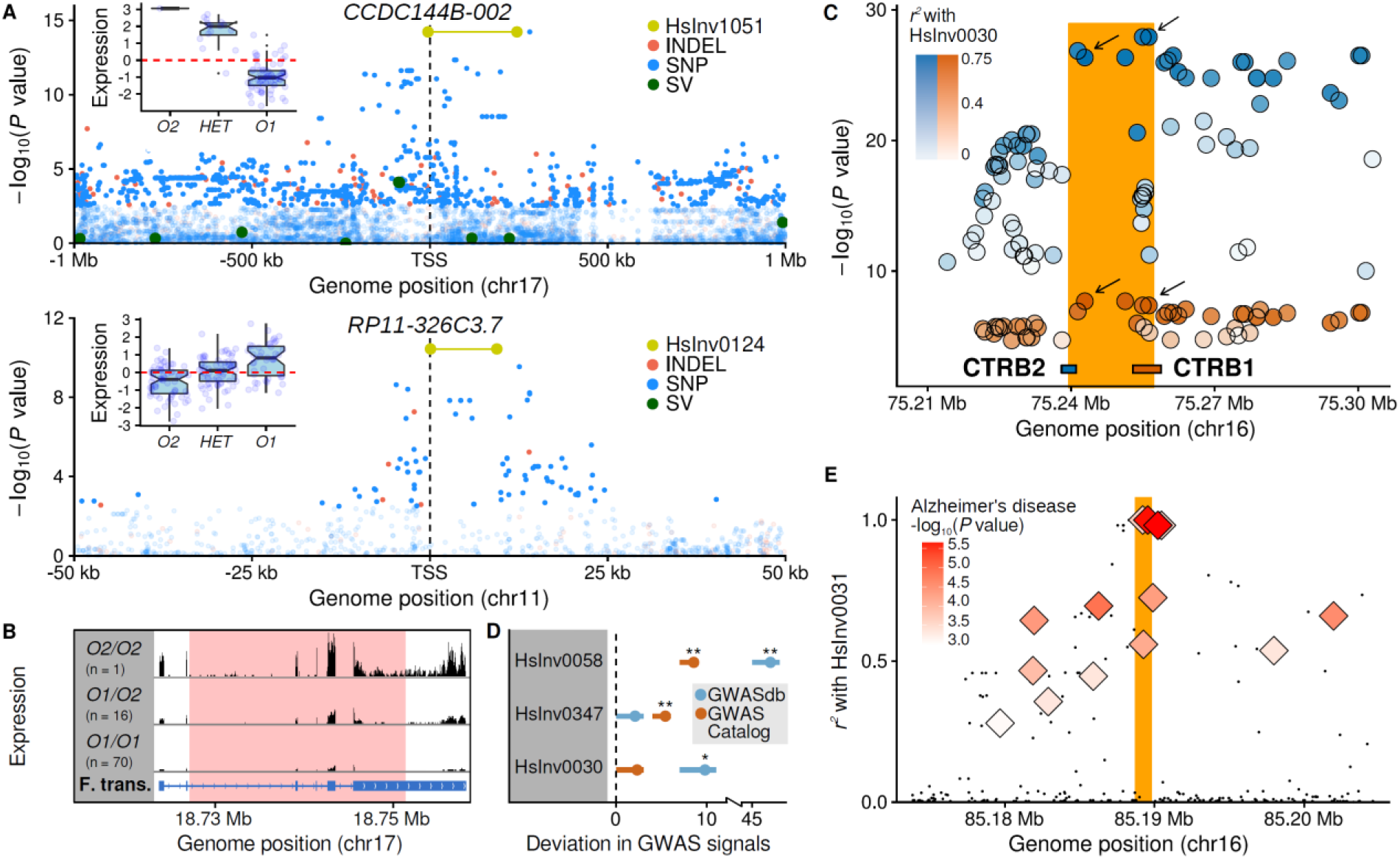
Examples of inversion functional effects. A. Manhattan plots for *cis*-eQTL associations in LCLs of transcript *CCDC144B-002* and gene *RP11-326C3.7*, showing inversions HsInv1051 (top) and HsInv0124 (bottom) as lead variants, together with boxplots of rank normalized expression and inversion genotype. B. Schematic diagram of average RNA-Seq expression profile in HsInv1051 genotypes from GEUVADIS LCL reads mapped to the inverted allele, showing the generation of a new fusion transcript. Inversion breakpoint is indicated in pink. C. Manhattan plot of pancreas GTEx-eQTLs for *CTRB1* (red) and *CTRB2* (blue) mapping around the HsInv0030 region (orange bar). Top lead eQTLs for each gene were the variants in highest LD with the inversion (rs9928842, *r*^2^ = 0.75; rs8057145, *r*^2^ = 0.73; black arrows). D. GWAS Catalog (orange) and GWASdb (blue) enrichment within individual inversions and flanking regions (± 20 kb) measured by the deviation in observed minus expected GWAS signals. Horizontal bars represent 95% one-sided confidence interval (onetailed permutation test: **, *P* < 0.01; *, *P* < 0.05). E. Linkage disequilibrium (LD) between HsInv0031 (orange bar) and 1000GP variants (black points) or Alzheimer’s disease GWAS signals (diamonds) (Pérez-Palma et al., 2014).

#### Inversions and phenotypic traits

We also investigated the role of inversions in phenotypic variation using genome-wide association studies (GWAS) collected in the GWAS Catalog (MacArthur et al., 2017) and GWASdb (Li et al., 2016) databases. First, we tested whether there was an enrichment of trait-associated signals in the inversion and flanking regions (±20 kb). We found a 1.26- and 1.95-fold increase of GWAS Catalog and GWASdb variants in inversion regions. When we looked for specific inversions driving this result (Figure S8A-B), the top one was the MHC-associated inversion HsInv0058, but HsInv0030 and HsInv0347 showed similar enrichment of GWAS hits in both datasets and significant differences from the expected number in at least one (Figure 5D). GWAS signals surrounding HsInv0030 are consistent with the genes it affects and involve type 1 and 2 diabetes, pancreatic cancer, insulin secretion, and cholesterol and triglyceride levels (Li et al., 2016; MacArthur et al., 2017). HsInv0347 is enriched in risk variants linked to glaucoma and optic disc and nerve characteristics (Li et al., 2016; MacArthur et al., 2017), which could be related to its effects in different tissues on the expression of *c14orf39* (*SIX6OS1*) and *SIX6*, related to eye development and located 92 kb away.

We also explored whether inversions were in strong LD (*r*^2^ ≥ 0.8) with known GWAS hits in the population or continent of the corresponding study (Table S12). That is the case of HsInv0004, which is in perfect LD in Europeans with an almost genome-wide significant GWAS SNP (*P* = 3.0 × 10^−7^), located ~24 kb away and related to asthma susceptibility in children (Sleiman et al., 2010), and with another one associated with body mass index in asthmatic children (P = 2.9 × 10^−4^) (Melén et al., 2013). Moreover, HsInv0006 is linked to diseases such as schizophrenia in Ashkenazi Jews (P = 5.0 × 10^−7^) (Goes et al., 2015) and glaucoma in Europeans (P = 2.7 × 10^−5^) (Gibson et al., 2012). Other inversions linked to trait-associated variants with lower significance levels (P = 10^−4^-10^−6^) include HsInv0045 and HsInv1053 with metabolic traits in urine (Suhre et al., 2011), HsInv0098 with coronary artery disease and cardiovascular prevention side effects (Isackson et al., 2011; Samani et al., 2007), and HsInv0409 with amyotrophic lateral sclerosis (Schymick et al., 2007). Remarkably, several of these inversions affect gene expression changes as well (Figure 2). A good example is HsInv0031, where the inverted orientation is associated with decreased expression of *FAM92B* in cerebellum and in almost perfect LD in Europeans (r^2^ = 0.98) with SNP rs2937145 associated with risk for Alzheimer’s disease (P = 2.02 × 10^−6^) (Pérez-Palma et al., 2014) (Figure 5E). Thus, these results indicate that inversions could have important phenotypic consequences. Nevertheless, for many inversions the low LD with SNPs means that their effects have been missed in typical array-based GWAS (Figure S8C).

#### Integrative analysis of genomic impact of inversions

In total, 9 inversions show strong signatures and 14 weak signatures of selection (of which 14 and 9 are likely due to positive and balancing selection, respectively) (Figure 2). Also, 22 inversions have potential functional consequences, ranging from altering the sequence of genes or non-coding transcripts (6), gene-expression changes in LCLs (8) or other tissues (14), and possible association with phenotypic traits (10) (Figure 2). More importantly, although not all functional and selection analysis could be applied to every inversion due to the lack of tag SNPs for many of them, we observed a significant enrichment of inversions with both effects on genes or gene expression and F_ST_ and LSFS selection signals directly linked to the inversion (Fisher’s exact test *P* = 0.0320) (Figure 2). This association is still significant when we consider only the strongest functional effects and selection signals (Fisher’s exact test *P* = 0.0130) or just the 21 inversions with perfect tag SNPs that were included in most analyses (Fisher’s exact test *P* = 0.0300). Therefore, both types of evidence independently support the important impact of inversions in the human genome.

One particularly interesting example is HsInv0201, an old inversion (>1.5 Mya) present also in Neanderthals and Denisovans (Table S7), which has an intermediate frequency around the globe and clear signals of balancing selection according to the LSFS and NCD1 tests (Figure 6A). This inversion deletes an exon of *SPINK14* and is associated with expression changes in several nearby genes, showing the highest LD with top eQTLs for *SPINK13* and *SCGB3A2* in the thyroid (Table S11). By analyzing additional expression data during immune response to pathogen infection from two independent studies (Alasoo et al., 2018; Nédélec et al., 2016), we found that the inversion is lead eQTL of *SPINK6* in Salmonella-infected cells (Figure 6B). In fact, the inversion deletes the promoter and first exon of a putative novel *SPINK6* isoform of this gene (Figure 6B), which may explain the lower expression levels of the gene associated with the inversion during infection. Moreover, the inversion is in high LD (*r*^2^ = 0.971 in EUR) with a secondary signal associated with the plasma levels of SPINK6 protein (Sun et al., 2018). Although the precise inversion consequences are unknown, the expression changes during infection, together with the role of several of the affected genes in lung and extracellular mucosas, suggest that they could be related to immune response, which is commonly under balancing selection (Bitarello et al., 2018).

In contrast, HsInv0006 is also a relatively old inversion (~450 kya) present in Neanderthals, in which the derived allele is close to fixation in many African populations but has intermediate frequencies in the rest of the world (Figure 6A). This particular distribution pattern, the high F_ST_ values and the LSFS test results all point to positive selection in Africa. Furthermore, the inversion is located within the first intron of the *DYSTK* gene (Vicente-Salvador et al., 2017) and is associated with expression changes in the proximal genes, including *DSTYK* upregulation in different tissues (Figure 6C; Figure S5). Interestingly, *DSTYK* deletion has been shown to cause pigmentation problems and elevated cell death after ultraviolet irradiation (Lee et al., 2017). Therefore, positive selection acting on these traits could explain the inversion frequency increase in Africa. Incidentally, the inverted orientation has been linked to higher risk of glaucoma in Europeans (Table S12) and glaucoma is more common and its effects more severe in individuals from African ancestry (Leske et al., 1994; Tielsch et al., 1991), supporting the functional relevance of the inversion.

Finally, there are several other interesting candidates that deserve further study (Figure 2). For example, HsInv0031 is the oldest inversion (>2 Mya), has global positive selection signals in the LSFS test, affects *FAM92B* expression in cerebellum, and has been associated with Alzheimer’s disease, which as other late-onset diseases could be influenced by pleiotropic effects of genes positively selected during human evolution (Corbett et al., 2018). Another inversion potentially related to immune response is HsInv0124, which shows frequency differences in Europe and affects several genes involved in virus infection defense, like *IFITM3* that is associated to reduced severity of flu infection in humans (Allen et al., 2017; Everitt et al., 2012; Wang et al., 2014; Zhang et al., 2013). HsInv0340 is found almost exclusively in Africa (MAF = 52.4%), and inactivates the lncRNA *LINC00395*, whose function is unknown but it is expressed at low levels in testis. Similarly, HsInv0059 may have been positively selected in Asian populations according to its higher frequency and the LSFS test, and is associated with expression changes in the gene within which it is located (*GABBR1*) plus the flanking gene (*PM20D2*). In the case of HsInv0389, it is almost fixed in Africa but has intermediate frequencies in other populations and is linked to expression differences in many genes (Figure S5). In addition, the region has been recurrently inverted multiple times in mammals, which may be due to a random neutral mutation process or some form of selection on the two alternative orientations (Cáceres et al., 2007).

**Figure 6.**
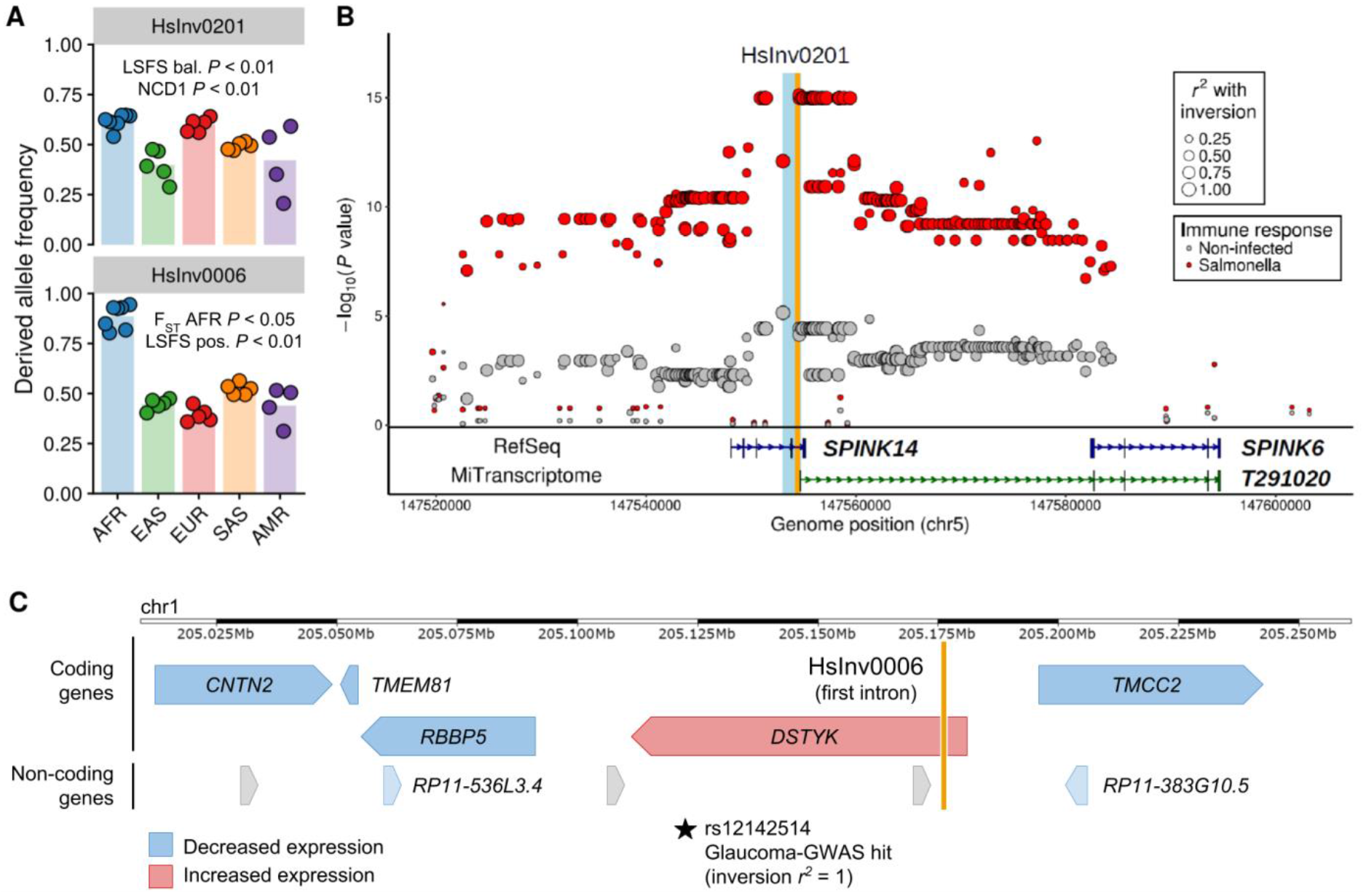
Integrative evolutionary and functional analysis of inversion candidates. **A**. Frequency of HsInv0201 and HsInv0006 across worldwide human populations from 1000GP Ph3 (colored dots) and in each population group (colored bars) estimated from global inversion tag SNPs (rs200056603 and rs79619752, respectively), showing a summary of selection test results. B. Manhattan plot for *cis*-eQTL associations with expression variation of gene *SPINK6* in infected and non-infected primary macrophages (Nédélec et al., 2016), showing SNPs in perfect LD with HsInv0201 (orange bar) as lead eQTLs in Salmonella infection. Variants are represented as circles with varying size depending on the LD with the inversion. The location of the genes in the region is shown below, including a new *SPINK6* isoform discovered by MiTranscriptome assembly (Iyer et al., 2015), whose first exon is removed by a deletion associated to the inversion breakpoints (light blue bars) C. Diagram of HsInv0006 (orange bar) genomic region showing the effect of the inverted allele on the expression of neighboring genes in different tissues according to the GTEx data and the inversion tag SNP in Europeans associated to increased risk of Glaucoma (Gibson et al., 2012). Arrowheads indicate the direction of transcription of the genes.

## DISCUSSION

This work represents the most complete and accurate study of human inversions so far, including a significant fraction of common inversions in the human genome (Martínez-Fundichely et al., 2014). Moreover, it is focused especially on inversions mediated by IRs, which escape detection by most techniques (Huddleston et al., 2017; Lucas-Lledó and Cáceres, 2013; Vicente-Salvador et al., 2017). As an example, less than one third of our inversions were detected in the recent description of 1000GP SVs (Sudmant et al., 2015), and they showed very high genotype error rates. In many cases this has precluded identifying tag variants that could be used to infer their potential effects (Chiang et al., 2017; Sudmant et al., 2015). Thus, the generation of reliable genotypes in multiple individuals of different populations has made possible to determine the functional and evolutionary impact of the great majority of the studied inversions for the first time.

In particular, we found clearly contrasting mutational patterns for inversions generated by homologous and non-homologous mechanisms, supporting a high-degree of recurrence of all inversions mediated by highly-identical IRs except three (two of which have very low frequency). Also, recurrence is not limited to humans but extends to other great ape species. Thanks to the additional number of individuals and populations analyzed, we have detected many recurrence events missed in small-scale studies (Aguado et al., 2014; Vicente-Salvador et al., 2017). By including multiple controls (such as PCR confirmation of genotypes in both breakpoints and different sources of nucleotide variation data) and a more detailed analysis of haplotype relationships, we have obtained a better estimate of the independent inversion events. This suggests that, like other repeats, IRs are rearrangement hotspots and the genome is more dynamic than previously thought. However, so far recurrence can be detected only through the sequences associated to the inversion, which limits its quantification, and we need more direct ways to determine inversion generation rates precisely.

Finally, similar to other types of potentially recurrent changes (Cantsilieris et al., 2018; Gymrek et al., 2016; Handsaker et al., 2015), the lack of association between SNPs and many inversions (along with poor coverage by common arrays in other cases) means that their effects on phenotypic traits have been largely missed in GWAS. As an example, half of the NAHR inversions cannot be predicted accurately with the most common imputation algorithms (Figure S4A). Our integrative analysis combining selection and function indicates that, as in other organisms (Jones et al., 2012; Joron et al., 2011; Küpper et al., 2016; Lowry and Willis, 2010; Thomas et al., 2008; Wellenreuther and Bernatchez, 2018), inversions can have important consequences in humans. In fact, inversions differ from other genetic variant because of their expected negative effect in fertility (Hoffmann and Rieseberg, 2008; Kirkpatrick, 2010), which is exemplified by the observed reduction in frequency with genetic length and the small number of inversions described compared to CNVs (Martínez-Fundichely et al., 2014). According to this, there should be a maximum length for an inversion to behave neutrally in terms of its effect on fertility, which could be as small as 1 kb. Above that size, some type of compensatory selection, perhaps related to advantageous regulatory changes on nearby genes, will be necessary for the inversions to reach a certain frequency. Therefore, this may explain the observed enrichment of inversions with functional and selection signals. Future in-depth studies of the identified candidates and other inversions will help uncover their real role in human evolution. Furthermore, the new developed inversion genotyping method will be instrumental to investigate their contribution to the unexplained part of the genetic basis of complex traits.

## ACKNOWLEDGEMENTS

We thank Cristina Aguado, Olga Dolgova, Teresa Soos, Esteban Urrea, David Vicente and Roser Zaurin for help with inversion genotyping, Antonio Barbadilla, Ruth Gómez, Óscar Lao, José Ignacio Lucas-Lledó, Sebastian Ramos-Onsins and Alfredo Ruiz for help with the evolutionary analysis, Mariona Bellet, Robert Castelo, Diego Garrido, Roderic Guigó and Rory Johnson for help with the gene expression analysis, Sònia Casillas and Alexander Martínez-Fundichelly for help with the inversion selection, and Xavier Estivill, Tomàs Marquès-Bonet, Aurora Ruiz-Herrera, the Coriell Institute for Medical Research, the Barcelona Zoo and the Banc de Teixits Animals de Catalunya (BTAC) for providing the human and non-human primate samples used in this study. This work was supported by research grants ERC Starting Grant 243212 (INVFEST) from the European Research Council under the European Union Seventh Research Framework Programme (FP7), BFU2013-42649-P and BFU2016-77244-R funded by the Agencia Estatal de Investigación (AEI, Spain) and the European Regional Development Fund (FEDER, EU), and 2014-SGR-1346 and 2017-SGR-1379 from the Generalitat de Catalunya (Spain) to MC, a PIF PhD fellowship from the Universitat Autònoma de Barcelona (Spain) to CGD, a La Caixa Doctoral fellowship to JLJ, and a FPI PhD fellowship from the Ministerio de Economía y Competitividad (Spain) to MO and IN. MGV was supported in part by POCI-01-0145-FEDER-006821 funded through the Operational Programme for Competitiveness Factors (COMPETE, EU) and UID/BIA/50027/2013 from the Foundation for Science and Technology (FCT, Portugal).

## AUTHOR’S CONTRIBUTIONS

MC conceived the inversion genotyping strategy, devised the study and oversaw all the steps; SV, MP and MC designed the genotyping assays; SV, DI, AD and MP carried out experiments; CGD, MGV, IO, CLF, MP and MC analyzed evolutionary data; CGD, JLJ, DC, BB, IN, PO, AA and LF performed selection tests; JLJ, MO, LP, TE and MC analyzed funcional effects; AB provided samples; MC, CGD, JLJ, MGV, LF and MP wrote the paper and all the authors contributed comments to the final version of the manuscript.

## ONLINE METHODS & SUPPLEMENTARY INFORMATION

### Studied inversions summary

This study is focused on 45 known polymorphic inversions from the InvFEST database (Martínez-Fundichely et al., 2014), which constitute a representative sample of human inversions and comprise most of those experimentally validated when the project started. We actually tried to include as many inversions as possible and the main limitation was the design of high-throughput genotyping assays, which prevented the analysis of those with large inverted repeats (IRs) (>25 kb) or for which there were no appropriate restriction enzymes cutting at each side of the IRs but not inside. The inversions were originally predicted in two main studies: a comparison of the independently assembled hg18 and HuRef genomes (Levy et al., 2007; Vicente-Salvador et al., 2017) and a fosmid paired-end mapping (PEM) survey of nine individuals (eight female and one male) with African (4 YRI), East Asian (1 CHB and 1 JPT) and European ancestry (2 CEU and presumably European individual NA15510) (Kidd et al., 2008; Korbel et al., 2007). Six inversions were detected exclusively in the genome comparison, 29 in the fosmid PEM analysis, and 10 by both methods (Table S1).

The 45 inversions had already been shown to be polymorphic in different studies by targeted PCR-based experiments (Martínez-Fundichely et al., 2014), including their genotyping in the validation panel formed by the nine individuals used in the fosmid PEM (Kidd et al., 2008). In addition, detailed breakpoint annotations in the hg18 reference genome and other inversion characteristics were already available from the existing studies, which were revised and updated here (Table S1). When necessary, the UCSC liftOver tool (Kent et al., 2002) was used to convert hg18 coordinates into hg19 or hg38. To avoid possible confusion about the ancestral status, the two inversion orientations have been named as orientation 1 (O1) and orientation 2 (O2). *O1* (AB and CD breakpoints) refers always to that found in hg18 and previously labeled as standard (*Std*), whereas *O2* refers to the alternative orientation (AC and BD breakpoints) previously labeled as inverted (*Inv*) (Martínez-Fundichely et al., 2014. The comparison with the 786 inversions predicted in the 1000 Genomes Project (1000GP) structural variants release (Sudmant et al., 2015) was done by extending the 1000GP coordinates by kb in each direction and using the breakpoint match batch query in the InvFEST database (Martínez-Fundichely et al., 2014). Only those cases in which there was overlap in both breakpoints, the inversion was considered to be the same.

### Human and ape DNA samples

We used a total of 551 human samples that belong to seven populations of four main population groups: Africa (AFR) (100 YRI and 90 LWK), Europe (EUR) (90 CEU, 90 TSI and individual NA15510), South-Asia (SAS) (90 GIH) and East Asia (EAS) (45 CHB and 45 JPT) (see Table S2 for details). Most individuals (481) do not share recent ancestors among them, whereas the other 70 are either children of mother-father-child trios (30 in YRI and 30 in CEU) or other first and second-degree relatives estimated based on sequence data (9 in LWK and 1 in GIH) (The 1000 Genomes Project Consortium, 2012, 2015). With the exception of NA15510, all individuals were included in the last phase of the HapMap Project (The International HapMap 3 Consortium, 2010), whereas 340 and 434 individuals were respectively included in phases 1 (Ph1) and 3 (Ph3) of the 1000GP (The 1000 Genomes Project Consortium, 2012, 2015) (Table S2). Genomic DNA samples come from lymphoblastoid cell lines (LCLs) commercialized by the Coriell Cell Repository (Camden, NJ, USA). In the case of 70 of the CEU samples plus 10 from YRI and NA15510, DNAs were extracted at the laboratory from LCLs as previously described, while the rest of CEU DNAs were acquired directed from Coriell (Aguado et al., 2014; Puig et al., 2015b; Vicente-Salvador et al., 2017).

Chimpanzee and gorilla samples include six DNAs extracted from frontal cortex tissue samples of the Banc de Teixits Animals de Catalunya (BTAC, Bellaterra, Barcelona, Spain) (N457/03, Z01/03 and Z02/03) or LCLs from individuals of the Barcelona Zoo (PTR1211, PTR1213 and PTR1215), which had been already used to genotype most of the inversions (Aguado et al., 2014; Vicente-Salvador et al., 2017). Moreover, we added 19 chimpanzee and five gorilla new DNA samples obtained from a previously-existing collection of primate LCLs (Table S6). These included four mother-father-child trios and one father-son pair in chimpanzees and one father-son pair in gorillas. As in the case of humans, cells were grown in 75-ml flasks to nearly confluency according to the recommended procedures and high-molecular weight genomic DNA was extracted using a standard phenol-chloroform extraction protocol (Aguado et al., 2014). All procedures that involved the use of human and non-human primate samples were approved by the Animal and Human Experimentation Ethics Committee (CEEAH) of the Universitat Autònoma de Barcelona.

### Experimental genotyping of inversions

As summarized in Figure 1, generation of the final inversion genotype data in humans and nonhuman primates was done in several steps: (1) high-throughput inversion genotyping with new MLPA-based assays; (2) quality controls to ensure genotype accuracy; and (3) validation of multiple genotypes by targeted PCR or iPCR.

Initial genotypes for all inversions, except four, were obtained by newly developed assays based on the multiplex ligation-dependent probe amplification (MLPA) technique (Schouten et al., 2002). MLPA consists on the simultaneous amplification of fragments of different sizes with common primers that are fluorescently labeled, and their detection by capillary electrophoresis. Each fragment is formed by a pair of oligonucleotide probes (left and right probes), which hybridize contiguously to specific genome regions in order to be ligated together in a subsequent step, and that include a variable stuffer sequence and the sequence of the forward or reverse common primers for multiplex amplification. This allows for a very specific detection of the region of interest and has been widely used for genome copy-number analysis. Here, we have applied a slightly-modified strategy for inversion genotyping by using two pairs of probes that bind specifically at the breakpoint sequences of each orientation (AB and CD or AC and BD), with one of the probes that could be common to both pairs (Figure 1). For inversions with IRs or other repetitive sequences at the breakpoints, we have developed a new method that we termed inverse MLPA (iMLPA) and uses a combination of inverse PCR (iPCR) and MLPA. To obtain an orientation-specific unique target sequence for these inversions, some extra processing is necessary before probe hybridization, which involves digestion with a restriction enzyme that cuts at each side of the repetitive sequences at the breakpoints followed by self-circularization of the digested DNA molecules by ligation in diluted conditions. Then, MLPA can be carried out as usual with a pair of probes that bind specifically at the self-ligation site of the circular molecules from each orientation (Figure 1). In total, 17 inversions were interrogated simultaneously with regular MLPA and 24 with iMLPA (Table S1). MLPA (68) and iMLPA (87) probes were designed using the program Proseek (Pantano et al., 2008) and manually modified to hybridize around the inversion breakpoints (MLPA) or the self-ligated restriction-enzyme target site (iMLPA) sequences, taking into account the usual MLPA-probe recommendations (Schouten et al., 2002) and that no common SNPs were close to the ligation ends of the probes (see Table S13 for probe sequences).

As mentioned before, iMLPA differs from normal MLPA by the addition of several extra initial steps and protocol optimization was carried out by comparison with the known genotypes from the validation panel of nine individuals (Martínez-Fundichely et al., 2014). A detailed description of iMLPA steps can be found in the patent application EP13382296.5 (Cáceres et al., 2015). Briefly, 400 ng of genomic DNA of each sample were first digested overnight at 37ºC in six separated 20-μl reactions with 5 U of the appropriate restriction enzyme (*EcoRl, Hindlll, Sacl* or *BamHI* from Roche and *Nsil* or *Bglll* from New England Biolabs), followed by restriction enzyme inactivation for 15 min at 65ºC (with the exception of *BglII* that was inactivated for 20 min at 80ºC). Next, DNA self-ligation was performed overnight for 16-18 hours at 16°C by mixing together the six restriction enzyme digestions with 1x ligation buffer and 400 U of T4 DNA Ligase (New England Biolabs) in a total volume of 640 μl (resulting in an optimal concentration of 0.625 ng/μl of the DNA fragments generated by each restriction enzyme). Then, the ligase was inactivated and the DNA was fragmented by heating at 95ºC for 5 min, and the DNA was purified with the ZR-96 DNA Clean & Concentrator™-5 kit (Zymo Research) and resuspended in 7.5 μl of water.

The last step was the regular MLPA assay using the SALSA MLPA kit (MRC-Holland), according to the manufacturer instructions with minor modifications. In particular, the ligated DNA was denaturated at 98ºC for 1.5 min, and probe hybridization was carried out adding 1.5 μl of our iMLPA probe mix (Table S13) plus 1.5 μl of SALSA MLPA Buffer (MRC-Holland) and incubating for 1.5 min at 95ºC and 16 hours at 60ºC. Ligation of adjacent probes was then performed for 25 min at 54ºC by adding 25 μl H_2_O, 1 μl SALSA Ligase 65, 3 μl Ligase Buffer A and 3 μl Ligase Buffer B (MRC-Holland). After ligase inactivation (5 min at 95ºC), PCR amplification of ligated probes was performed separately in three groups of 8-9 inversions (Table S13) using a common reverse primer and one of three forward primers labeled with a different fluorochrome in each case (FAM, VIC or NED) (Table S14). Each PCR was done in 25 μl with 5-6 μl of the iMLPA hybridization-ligation reaction, 2 μl SALSA PCR buffer (MRC-Holland), 0.25 mM each dNTP, 0.2 μM each primer, 1 μl PCR buffer without MgCl_2_ (Roche), and 2.5 U of Taq DNA polymerase (Roche). PCR conditions were 95°C for 15 sec, 47 cycles of 95°C for 30 sec, 60°C for 30 sec and 72°C for 60 sec, and final extension at 72°C for 25 min.

Finally, 0.67 μl of the three PCRs of each sample were mixed together, analyzed by capillary electrophoresis using an ABI PRISM 3130 Genetic Analyzer (Applied Biosystems), and the peaks were visually inspected using the GeneScan version 3.7 software (Applied Biosystems). For the regular MLPA, the process was identical with the exception that it started directly at the denaturation step of 100-150 ng of genomic DNA in 5 μl for 5 min at 98 °C and that the ligated probes were amplified in only two multiplex PCRs with 8-9 inversions each (Table S13). In both cases all the successive reactions were carried out in a 96-well plate format to maximize speed and throughput and, with the exception of those used for optimization of the technique, only one MLPA or iMLPA reaction was done for every sample.

In addition, we carried out PCRs and iPCRs to genotype seven inversions in the 551 human samples (three already included and four not included in the MLPA/iMLPA assays) and to validate multiple of the MLPA/iMLPA genotypes (Table S1). Multiplex PCRs and iPCRs of all the inversions had been previously set up and were done as described before (Aguado et al., 2014; Martínez-Fundichely et al.; Vicente-Salvador et al., 2017), using primers flanking either the breakpoint (PCR) or the self-ligation site of circularized molecules (iPCR) (Table S14). PCR/iPCR validations were done for those genotypes identified in the quality controls plus several additional samples chosen at random. Genotype quality controls included: (1) Comparison with available genotypes from the same individuals (Martínez-Fundichely et al., 2014); (2) Consistency in mother-father-child trios; (3) Association with other polymorphic variants and haplotypes (as explained in detail below); and (4) Detection of polymorphic restriction enzyme target sites that could affect the iPCR or iMLPA assays. In the restriction site polymorphism analysis, we downloaded all known SNPs and indels around the inversion region (±50 kb) from dbSNP version 142 and determined if the alternative sequence with each of the variants has the recognition sequence of the restriction enzyme used in the inversion genotyping assay. For all potential restriction site gains or losses affecting the iPCR/iMLPA experiments, we checked their frequency in 1000GP populations and we confirmed if they result in missing bands of any of the orientations in our genotyped individuals. To ensure that the genotypes were completely accurate, in most MLPA/iMLPA potential discrepancies and especially in those cases that could be only validated by iPCR, both breakpoints of each orientation were tested.

The work described here is based on version 4.8 of the human inversion genotype data, which includes the latest information from the PCR/iPCR validations, except for some of the first analyses, which use the previous version 4.7 and are specified in each case. The two datasets differ by the addition of 10 new genotypes and 11 genotype corrections, which affect in total 10 inversions. Inversion frequency was calculated with the 480 non-related individuals or the 434 individuals in common with 1000GP Ph3 depending on the analysis, and using either the minor allele frequency (MAF) or the derived allele frequency (DAF), as stated. The minor allele was defined based on the frequency in the seven populations combined. Deviation from Hardy-Weinberg equilibrium in each population was calculated with Plink—hardy option (Purcell et al., 2007) considering an exact test *P* < 0.01.

For inversion genotyping in chimpanzees and gorillas, first the available genome sequences of each species (panTro4, panTro5, gorGor4 and gorGor5) were examined for nucleotide differences that may affect the human assays (primer binding sites, gains or losses of restriction sites, etc.). Then, we ran the same MLPA and iMLPA protocols as described above. For those inversions where MLPA/iMLPA did not work, we obtained the genotypes by PCR or iPCR. While we could easily obtain results for virtually all NH inversions, some assays involving iPCR used to genotype inversions with IRs in humans (which require conservation not only of primers but of restriction sites) did not work in nonhuman primates. In some cases, chimpanzee or gorilla specific primers and even new restriction enzymes for iPCR were used to overcome the human assay problems (Aguado et al., 2014) (Table S6; Table S14). However, this was not possible for all inversions in both species due mainly to genome gaps, resulting in not testing a few inversions in one of the species and HsInv0055, HsInv0374 or HsInv1051 in any of them. Genotypes of parent-child trios and father-son pairs in chimpanzees and gorillas were consistent with expected genetic transmission in each inversion. In addition, all polymorphic inversions in chimpanzees or gorillas were validated by PCR or iPCR to make sure that there were no errors in the iMLPA results (and whenever possible the two breakpoints were interrogated). The analyzed breakpoints and assays used are summarized in Table S6, which include 40 inversions in chimpanzee (16 MLPA, 5 iMLPA, 3 PCR, 11 iPCR, 1 MLPA + PCR, and 4 iMLPA + iPCR) and 41 inversions in gorilla (15 MLPA, 4 iMLPA, 6 PCR, 11 iPCR, 1 MLPA + PCR, and 4 iMLPA + iPCR). Of those, 26 inversions were tested only by one breakpoint and 14 by both breakpoints in chimpanzees, whereas 30 inversions were tested only by one breakpoint and 11 by both breakpoints in gorillas. Nevertheless, the primate genotyping results should be taken with caution, because despite the assay checks, unaccounted polymorphisms may prevent the detection of one of the alleles by iPCR and lead to incorrect genotypes, especially when only one breakpoint can be interrogated.

### Analysis of nucleotide variants associated with inversions

We measured the correlation between inversion genotypes and the genotypes of overlapping and neighboring biallelic variants (up to 200 kb at each side of the inversion) in two complementary data sets: 1000GP Ph3 (including SNPs and indels for 434 unrelated individuals) and HapMap phase 3 (that includes fewer SNPs but all the 480 unrelated individuals). Pairwise linkage disequilibrium (LD) between variants was calculated by the *r*^2^ statistic for each population independently, as well as by population group and globally with all individuals together. Variants located within the breakpoint interval and associated deletions or IRs were excluded to avoid possible genotyping errors. LD in 1000GP data was analyzed with plink v1.90 (Purcell et al., 2007) using the available BCF files (ftp://ftp.1000genomes.ebi.ac.uk/vol1/ftp/release/20130502/). In the case of HapMap variants (HapMap_release27), LD was calculated with Haploview v4.1 (Barrett et al., 2005) and since the set of SNPs genotyped in each population was different, LD was calculated for all SNPs available in each population separately and for those tested in all populations.

Overall, we found that 23 inversions have at least one perfect tag variant (*r*^2^ = 1) in the global sample, including all NH inversions except one (HsInv0102). Consistent results were obtained in 1000GP and HapMap data, with 27/29 of the 1000GP perfect tag SNPs tested in HapMap having also perfect LD with the inversion. In addition, HapMap data adds three perfect tag SNPs in the list that are either in high LD or were not reported in 1000GP. Apart from the clear contrasting pattern between NH and NAHR inversions, there was considerable variation in the number of tag variants between inversions, which is probably related to the recombination events outside the inverted region. As expected, the number of inversions with tag variants (*r*^2^ > 0.8) increases when focusing on individual populations or populations groups, possibly reflecting population-specific haplotypes associated with the inversion and for all except five NAHR inversions there are some variants with *r*^2^ > 0.8 in at least one population. This analysis allowed us to detect specific individuals in a few inversions whose genotype did not match that expected from the tag SNPs. Therefore, they were checked independently by PCR or iPCR and most of them were confirmed to be genotyping errors (Figure S2).

SNPs and indels around inversion regions were further classified according to its distribution across orientations using in-house perl and bash scripts that take into account the genotypes of heterozygotes and both homozygotes for the inversion, as previously described (Aguado et al., 2014; Vicente-Salvador et al., 2017). The classification was done as follows: if a variant is unambiguously polymorphic in the two orientations, it is classified as shared; if it is clearly polymorphic only in one, it is classified as private; if it is monomorphic in both orientations and they have different alleles, it is classified as fixed (corresponding to those in perfect LD with the inversion). To minimize possible genotyping errors in 1000GP data, only the most reliable variants according to the 1000GP strict accessibility mask were included. As mentioned in the Results, most NH inversions have no shared variants between orientations within the inverted region, consistent with a complete inhibition of recombination in inversion heterozygotes, although we could only test 13 NH inversions with accessible variants inside. The only exceptions were HsInv1053, that has a single shared SNP near the second breakpoint in 1000GP and few shared SNPs in HapMap, and HsInv0095, with shared variants only in HapMap. These variants tend to be grouped in certain positions and are likely the result of gene conversion, SNP genotyping errors or even independent mutations. Conversely, shared SNPs of inversions mediated by NAHR tend to be scattered along the inverted region (Figure 3B).

The distribution of fixed and shared variants obtained using the 1000GP data (which has more resolution than HapMap) allowed us to define a region surrounding the inverted region with little or no recombination between the two orientations. This extended flanking sequence is useful to increase the number of informative sites to understand inversion evolutionary history and detect possible selection signals, especially for very small inversions. Non-recombining flanking regions were defined as the longest sequence at each side of the inversion breakpoints up to a maximum distance of 20 kb that: (1) does not contain reliable shared variants compatible with a crossing-over event between *O1* and *O2* chromosomes; and (2) includes most of the fixed variants (Table S3). For NAHR inversions with a high proportion of shared variants within the inverted region and no perfect tag variants, no extra flanking region was considered.

### Phasing of inversions and haplotype visualization

For some analyses it is necessary to determine the sequence of the chromosomes with each orientation. Since this could be affected by different sources of error, we followed two complementary strategies to minimize phasing errors and obtain more robust results. First, after testing several common phasing programs, we inferred the phase of nucleotide variants and inversion alleles using PHASE 2.1 (Stephens and Donnelly, 2003; Stephens et al., 2001). One advantage of PHASE is that it is possible to fix the phase of the two markers representing the inversion orientation that were added at the position of the breakpoints, avoiding switch errors in heterozygotes (Aguado et al., 2014; Vicente-Salvador et al., 2017). Phasing was done independently for each population group (EUR, SAS and EAS), except for the two African populations (YRI and LWK), which were phased separately, using the available trio information when possible. We performed five iterations (-x5) for HapMap data and two iterations (-x2) with the hybrid model (–MQ) for 1000GP data. Due to the computing time requirements, we selected only the variants within the inversion region (eliminating the breakpoint intervals and any associated deletions or IRs, as before) plus 20 kb of flanking sequence at either side for 1000GP Ph1 data (which was the only one available at that time) and 200 kb for HapMap data (to ensure the inclusion of enough polymorphic sites). Second, we took advantage of the already available phased haplotypes from the 1000GP Ph3 data, which have been shown to be very accurate (The 1000 Genomes Project Consortium, 2015). In this case, for those inversions with perfect tag variants (*r*^2^ = 1) the inversion orientation of the haplotypes was imputed based on the presence of the linked variant alleles. As expected, the alleles of the different variants associated to a given orientation were always found together in the haplotypes. For the rest of inversions without perfect tag variants, only homozygotes or hemizygotes for each orientation were analyzed, in which the inversion status of the haplotypes can be assigned unambiguously.

Next, we tried to reconstruct and visualize the relationships between the different haplotypes for each dataset. Recombination between the sequences complicates the interpretation, especially for large regions. Thus, we combined several data sources and representation methods. First, we used the phased 1000GP Ph1 haplotypes for the inverted region to build Median-Joining (MJ) networks (Bandelt et al., 1999) with the NETWORK v.4.6.1.3 software (www.fluxus-engineering.com). The same analysis was also done for the HapMap phased haplotypes, including in this case part of the flanking region for small inversions, if needed. Reticulated networks are able to accommodate past recombination events, but each sequence is reduced to a node or edge, making it difficult to understand at the same time haplotype relationships and the spatial distribution of nucleotide changes along the sequence. Therefore, we devised our own way to represent the similarities between haplotypes, named integrated haplotype plot (iHPlot), which combines a hierarchical clustering, distance matrix and visual alignment of the alleles in each polymorphic position, plus the haplotype orientation and the populations in which is present (see Figure S3). These are similar to the Haplostrips plots that have been recently developed independently (Marnetto and Huerta-Sánchez, 2017). Specifically, distances between simplified haplotypes after removing singleton positions were computed as the number of pairwise differences and were clustered with the UPGMA average method implemented in R (R Core Team, 2017) base function hclust. The corresponding dendrogram was then created using ggdendro R package (de Vries and Ripley, 2016) and all the information was integrated with a custom R script using ggplot2 and cowplot packages (Wickham, 2009). iHPlots were applied to the phased 1000GP Ph1 haplotypes of the inverted region and the imputed 1000GP Ph3 haplotypes based on inversion tag SNPs or homozygotes of each orientation. For 1000GP Ph3 data, only accessible SNPs (excluding indels) according to the pilot accessibility mask, which includes more SNPs than the strict mask, were used. In addition, besides the inverted region, whenever possible, we extended the analysis to the non-recombining region flanking the breakpoints (excluding associated indels and IRs) to increase the resolution of haplotype discrimination.

To estimate more reliably inversion evolutionary history, we combined the information from the different strategies for phasing and visualization of the haplotypes of each region: 1) MJ networks of 1000GP Ph1 phased data; 2) iHPlots of 1000GP Ph1 phased data; and 3) iHPlots of 1000GP Ph3 published haplotypes (including when possible the flanking non-recombining region). Moreover, HapMap phased data was used to confirm 1000GP results, although in many cases information was available from just a few SNPs. For inversions with perfect tag variants (*r*^2^ = 1), results were based mainly in the extended 1000GP Ph3 haplotypes, which allowed the filtering of accessible SNPs and included less phasing errors due to the use of sequences from more individuals (The 1000 Genomes Project Consortium, 2015). All inversions could be analyzed except HsInv0041, which did not have enough variants and was excluded. Results were consistent with those of the other analyses, with the exception of a few phasing errors by PHASE in 1000GP Ph1 data. In addition, in HsInv0409 there appears to be an imputation error in *O2/O2* individual NA20530 in 1000GP Ph3 (in which one of the haplotypes is typical of *O1* chromosomes, whereas in 1000GP Ph1 both haplotypes belong clearly to the *O2* group).

For inversions without tag variants, we relied mainly in the phased 1000GP Ph1 haplotypes, since it was possible to analyze all genotyped individuals in common, although there could be more phasing errors in inversion heterozygotes. First, we defined the putative original inversion event based on the ancestral information, the haplotype diversity within each orientation, and the frequency and geographical distribution of haplotypes, tending to favor as the first event those occurring in Africa. Next, we conservatively identified potential additional inversion or re-inversion events in differentiated clusters of haplotypes with both orientations. In order to consider that there has been inversion recurrence, these clusters have to differ from all other ones, and especially from those with the same orientation as the potential recurrence event, by three or more sequence changes along most of the inverted region (and spanning at least 2 kb). Therefore, the presence of these nucleotide differences cannot be explained easily by other mechanisms, such as gene conversion or sequence errors. Direction of recurrence events was defined based on the relationship between the clusters and the frequency of the haplotypes with each orientation (Table S4). Possible phasing errors in inversion heterozygotes were checked manually by determining if switching the orientation of both haplotypes still supports unequivocally the existence of recurrence. In addition, the analysis was repeated in 1000GP Ph3 iHPlots in which just the orientation from haplotypes of *O1* and *O2* homozygotes is assigned, and only those clear recurrence events not invalidated with the new data were considered. Finally, inversion genotypes of 1-25 individuals supporting each of the independent recurrence events were validated by PCR or iPCR (in this case testing mostly both breakpoints) to discard possible genotyping errors (Table S4). Moreover, we validated the genotype of many more individuals with rare orientation-haplotype combinations of these inversions, plus of other inversions with a high proportion of shared SNPs for which recurrence events could not be clearly identified. The analysis was done with inversion genotype data version 4.7, and as previously mentioned the PCRs allowed us to detect 11 genotype errors that were updated in version 4.8 of the data. Of those, four correspond to potential recurrent individuals, but they did not invalidate any of the predicted recurrence events.

In the case of HsInv0832, we gathered the publicly available information of the chr. Y haplogroups of 232 of the 282 genotyped males from different sources (Van Geystelen et al., 2013; Hallast et al., 2015; Magoon et al., 2013; Poznik et al., 2016; Wang and Li, 2013; Wei et al., 2013), which is shown in Table S5. Most of these studies determined also the evolutionary relationship between the chr. Y haplogroups, which were largely consistent and are shown in a simplified genealogical tree using the branch lengths of Poznik *et al*. (2016) (Poznik et al., 2016) in Figure 3. This allowed us to identify with confidence five independent inversion events in the HsInv0832 region, assuming the most parsimonious scenario. HsInv0832 inversion rate was estimated dividing the number of inversion events (*n*) by the number of generations (*g*) encompassed in the phylogeny that relates the 217 Y-chromosomes for which sequence data was available (Poznik et al., 2016). To estimate *g*, we used the data from the B-T branch split to the leaves from Poznik *et al*. (2016) (Poznik et al., 2016), including a B-T branch split time of 105.8 kya, a total number of mutations of all branches involved in the phylogeny that relates those 217 males of 17,362, and an average number of mutations of all branches of 784.08, plus a generation time of 25 years as in Repping *et al*. (2006) (Repping et al., 2006). This results in 93,710 generations and an inversion rate of 5.34 × 10^−5^ per generation. In addition, to have another estimate of the inversion rate, we also used the approach of Hallast *et al*. (2013) (Hallast et al., 2013), which was based on Repping *et al*. (2006) (Repping et al., 2006) that resequenced 80 kb in 47 Y chromosomes covering most major branches of the phylogenetic tree to obtain the nucleotide divergence in an unbiased manner. Based on their data, we estimated a lower and upper bound of *g* of 127,467-336,533 generations by calculating the maximum (631) and minimum (239) total number of mutations spanning all the informative branches and an average number of mutations to the root in the different branches of 8.85, assuming a divergence time of 118 kya and a generation time of 25 years (Hallast et al., 2013; Repping et al., 2006). This yields an inversion rate of 1.48-3.92 × 10^−5^, which is quite similar to the previous one.

### Bioinformatic analysis of inversion orientation in other genomes

To complement the results from the experimental genotyping in chimpanzees and gorillas, the ancestral state of inversions was also estimated by checking their orientation in several of the best assemblies from non-human primate species already available: chimpanzee (panTro5), gorilla (gorGor5), orangutan (ponAbe2) and rhesus macaque (macRhe8). The analysis was done using an automated bash script based on the command-line blat tool (v35×1) (Kent et al., 2002). For each inversion, three separate hg18 sequences were extracted using twoBitToFa UCSC utility: the inverted region (or alternatively two separate internal 10-kb sequences adjacent to each breakpoint when the inverted region is longer than 20 kb) and the two 10-kb segments flanking each breakpoint outside the inversion. We excluded the breakpoint intervals and their associated IRs and indels to avoid ambiguous mappings. Then, each sequence was aligned with blat to the genomes of interest, which were downloaded from the UCSC Genome Browser website in 2bit format. The longest hit was kept as the likely homologous region in the target assembly and orientation was defined as *O1* if all best hits mapped in the same strand and as *O2* if the internal best hit(s) mapped in the opposite strand than those from the external sequences. As quality control, all best hits needed to be in the same scaffold or chromosome and the total span in the target assembly had to be 0.5-2-times that in hg18. In addition, results from the automated analysis were revised by aligning the sequences spanning the entire region from each assembly with the Gepard dotplot application (Krumsiek et al., 2007) and Blast2seq (Altschul et al., 1990), using default parameters, especially in those cases in which the orientation could not be reliably defined or was inconsistent across species or with published data (Aguado et al., 2014; Vicente-Salvador et al., 2017).

Moreover, we also checked for the presence of the inversions in several available ancient hominin genomes. Due to the fragmented status of these genomes, we used a different strategy based on the identification of reads that span the inversion breakpoints. In particular, we downloaded the raw reads of the Neanderthal (Prüfer et al., 2014), Denisovan (Meyer et al., 2012) and two ancient modern human genomes, Otzi (Keller et al., 2012) and La Braña 1 (Olalde et al., 2014), and the analysis was restricted to 19 NH inversions without IRs at the breakpoints. For each inversion, a breakpoint library with four 100-bp sequences centered at the two breakpoints of both orientations (Table S15) were mapped to the reads from each genome with a slightly modified version of the BreakSeq pipeline, as previously described (Lucas-Lledó et al., 2014; Puig et al., 2015b; Vicente-Salvador et al., 2017). In addition, in some cases in which BreakSeq did not identify any reads mapping to the inversion breakpoints, the possible genotype was inferred based on the visual inspection of the alignment of reads to the human genome. A particular orientation was considered to be reliably identified in a genome if it was supported by at least two reads from any breakpoint (Table S7).

### Inversion age estimate

Age of unique inversions was estimated with the usual divergence-based approach. The differentiation accumulated since the inversion origin is usually calculated as the mean pairwise nucleotide differences between sequences in opposite orientations after subtracting the average differences expected in the original population (without the inversion), which is typically approximated by the nucleotide variation within the ancestral orientation (Corbett-Detig and Hartl, 2012; Hasson and Eanes, 1996). Specifically, we estimated pairwise differences between orientations from the available 1000GP Ph3 haplotypes, taking all populations together and using all SNPs (excluding indels). To ensure the maximum reliability of the divergence estimates, as before, sequence orientation was determined by the presence of tag variants or by using only *O1* and *O2* homozygous individuals. In all cases, we considered the inverted region, plus the extra non-recombining flanking region if available (up to a maximum distance of 20 kb), which allowed us to have more information in short inversions. Breakpoint intervals including IRs or associated indels were excluded to avoid sequence errors. For two low-frequency inversions, HsInv0061 and HsInv1051, divergence could not be estimated because there were no tag variants in the analyzed region and all inversion carriers are heterozygous. In order to control for the noise in the nucleotide difference values, we repeated the estimates by bootstrap sampling the same number of total individuals with replacement 1000 times. A first age estimate was obtained by calculating the average differences in the original population as the largest of the average pairwise differences within *O1* or *O2* sequences and using a constant substitution rate of 10^−9^ changes per base-pair per year (Moorjani et al., 2016). Moreover, in order to control for local differences in substitution rates, we obtained two additional local estimates from the divergence with chimpanzee and gorilla genomes, considering, respectively, 6 and 8 million years as split times with humans (Table S7). Pairwise LASTZ alignments (Harris, 2007) of human hg19 assembly with chimpanzee (assembly CSAC 2.1.4/panTro4) and gorilla (assembly gorGor3.1) genomes were retrieved from ENSEMBL GRCh37 portal (Yates et al., 2016), using the Compara Perl API (Herrero et al., 2016). We then used Kimura’s two-parameter substitution model (Kimura, 1980) to calculate the divergence between human and outgroup assemblies in the same region analyzed above, after removing alignment gaps and non-syntenic alignment blocks. Alignments shorter than 1 kb were discarded, including chimpanzee alignment for inversion HsInv0045 and both outgroup alignments for inversion HsInv0041.

### Inversion frequency analyses

To measure and control the effect of the ascertainment bias associated to the study design in the observed frequency of inversions, we simulated the detection and genotyping process in biallelic SNPs from 1000GP Ph3, using a bash and R (R Core Team, 2017) pipeline. Given the heterogeneous origin of the inversions included in the study (Table S1), we simulated two different processes: one for the 38 autosomal or chr. X inversions detected from the fosmid PEM data of nine individuals (Kidd et al., 2008); and another for the six inversions detected exclusively by comparison of two genome assemblies (Levy et al., 2007). The process was simulated in two steps: (1) select those SNPs where the alternative allele is present in the detection panel; and (2) keep only the fraction of SNPs equivalent to the probably of detection of the inversions with the methods employed. In the first step, we built panels with matching demographic and gender composition to the detection samples from 1000GP individuals. For the PEM study, we were able to use all the original individuals except for NA19240 and NA15510, which were replaced with NA18502 and NA12717. For the genome comparison study, we used a randomly selected European male (NA12872) and selected all SNPs that contained the alternative allele. Variants were filtered from 1000GP vcf files (ftp://ftp.1000genomes.ebi.ac.uk/vol1/ftp/release/20130502) with bcftools v1.7 view command (Li, 2011. Additional filters were applied to the SNPs to simplify comparisons (keeping only SNPs with rsID and ancestral allele determined in 1000GP vcf files), to use only putatively neutral variants (conservation GERP score below two in functionally annotated 1000GP vcf files ftp://ftp-trace.ncbi.nih.gov/1000genomes/ftp/release/20130502/supporting/functional/annotation/unfiltered/), and to ensure high SNP quality (accessible according to the 1000GP Ph3 strict accessibility mask). However, the effect of these extra filters on the final frequency distribution was negligible, affecting just <1.5% of the detectable SNPs, which have similar average frequency to the rest of SNPs.

The second step was only simulated for the PEM data, since the limitations of inversion detection by assembly comparison are likely independent of variant frequency (and instead probably just related to repeat content and complexity of the genomic region). PEM detection, on the other hand, is affected by the sizes of the inversion and the IRs at the breakpoints, both of which limit the number of PEMs supporting it. To that end, we modeled the detection of an inversion that is present in the PEM panel as a function of these two characteristics and the number of chromosomes with the alternative orientation. Specifically, the probability of having two discordant PEMs in the whole panel (the minimum number necessary to detect an inversion) was calculated by a Poisson distribution with a lambda parameter equal to the expected number of discordant PEMs (E(*disc*)). Following Equation 1 in Lucas-Lledó *et al*. (2013) (Lucas-Lledó and Cáceres, 2013), E(*disc*) for the two breakpoints of a given inversion (*inv*) and IR (ir) size, considering the average PEM insert length (*ins*) and read length (*read*), was estimated as:

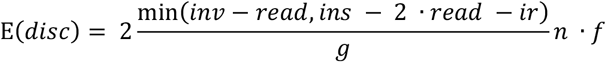

where *g* is the sequenced haploid genome size (approximated to 3 Gb), *n* is the total number of fosmids sequenced (Kidd et al., 2008; Tuzun et al., 2005), and *f* is the fraction of chromosomes carrying the mutation in the nine individuals analyzed. For each inversion, a custom R script was used to generate a matching random sample of 10,000 SNPs. These SNPs were selected from the total pool of polymorphic SNPs in the PEM panel according to its detection probability based on its frequency and the inversion characteristics, including the chromosome type (autosomes or chr. X).

The frequency distributions of the SNPs in the 434 1000GP Ph3 samples in common to our study were then compared with that of the inversions. Comparisons were done globally and by population group both for all inversions together and stratifying them by the two main generation mechanisms (using in each case the subset of SNPs matched to the inversions that are analyzed). Mean and median frequencies of inversions and SNPs were compared by sampling 10,000 sets of SNPs without replacement, with one matched-SNP per inversion at a time to preserve any differences between them. Empirical *P*-values were estimated as twice the fraction of samples with values more extreme or equal than the observed. Similar results were obtained by comparing the distributions with a Kolmogorov-Smirnov non-parametric test.

Frequency differences between populations were estimated using the F_ST_ statistic (Weir and Cockerham, 1984). To calculate F_ST_ we created vcf files containing the inversion genotypes for the 434 individuals common to 1000GP Ph3 and used—weir-fst-pop option from vcftools (v0.1.15). F_ST_ values were obtained for each pair of populations within the same population group, each pair of population groups, and globally. Empirical null distributions were obtained from all 1000GP Ph3 bialellic SNPs polymorphic in the same 434 individuals that are accessible according to the strict criteria and have a defined ancestral allele. In order to control for FST dependence on chromosome type and allele frequency, we divided the null distribution by chromosome type (autosome or chr. X) and in 10 global MAF bins ranging from 0 to 0.5 (all including a minimum of 10,000 SNPs). Empirical *P*-values for each of the comparisons were estimated as the fraction of SNPs in the distribution corresponding to the same global MAF bin as the inversion with equal or larger F_ST_ (Table S8). Inversions with F_ST_ values within the top 1% of the empirical distribution (P < 0.01) in at least one of the comparisons were considered as strong candidates to be under positive selection in the populations with higher frequency, and those in the top 5% (P < 0.05) were considered weak positive selection candidates. Reduced levels of population differentiation are sometimes interpreted as evidence of balancing selection. However, power to detect the extreme low F_ST_ values was very low. Global population differentiation for all inversions together was measured by a hierarchic analysis of molecular variance (AMOVA) according to geographic criteria using Arlequin v3.5 (Excoffier and Lischer, 2010). Resulting variation was mainly due to the difference between the three continental groups for both autosomal inversions (F_ST_ = 0.13, *P* < 0.0001; F_CT_ = 0.11, *P* < 0.0001; F_SC_ = 0.03, *P* = 0.02) and chr. X inversions (F_ST_ = 0.24, *P* < 0.0001; F_CT_ = 0.20, *P* < 0.0001; F_SC_ = 0.05, *P* < 0.0001).

### Genomic determinants of inversion frequency

To study the factors that could affect the frequency of human inversions we tested multiple different genomic variables: (1) physical length of the inversion (measured as the internal segment within the breakpoints); (2) genetic length of the inversion (in 4N_e_r units); (3) number of genes within the inversion or breakpoint regions; (4) distance to the closest coding gene; (5) total number of significantly constrained sites in mammalian phylogeny inside the inversion (Davydov et al., 2010); (6) functional impact according to the direct effect of inversions on genes (see Table S9); and (7) the size of the IRs at the breakpoints of some inversions. Inversion genetic length was estimated from the SNP data from the 434 samples present in 1000GP Ph3 using rhomap function of the LDhat v2.2 package (Auton and McVean, 2007), and the cumulative 4N_e_r for all the chromosomes was used as a proxy for the inversion genetic length. The analysis was done separately using the DAF for NH inversions (with known ancestral allele) and the MAF for NAHR inversions (with no clear ancestral allele in most cases). First, we modeled the frequency of autosomal and chr. X inversions in each population group by correlations with the above inversion sequence characteristics. Since inversion genetic and physical length, IR size, the number of genes within the inversion, and the inversion functional impact showed a high multicollinearity when two or more of these variables were included in a regression model (high variance inflation factor), we decided to build the model by stepwise forward selection of predictors, keeping only variables with significant regression coefficients to avoid overfitting the model. We found a statistically significant logarithmic decrease of frequency with genetic length for all inversions together, which was the main genomic determinant of frequency. In addition, MM-regression models were built in R using function lmrob from robustbase package (Maechler et al., 2018) to reduce the effect of potential outliers in a reduced sample set. For NH inversions, the log-transformed inversion genetic length is the best predictor of DAF (variance explained 23-55%) (Figure 4B), being slightly better than the log-transformed physical size alone (variance explained 19-54%). In order to control for the detection bias against low-frequency small inversions, we repeated the regressions excluding small inversions (<2 kb) with similar results (Figure 4B). In the case of NAHR inversions, the logarithmic relationship of MAF with inversion genetic length has a much weaker predictive power (variance explained 5-14%) than the ratio of IR to inversion physical size, which explains 13-45% of the variance and has a positive regression coefficient in all population groups (AFR: β = 0.18, *P* = 0.029; EUR: β = 0.15, *P* = 0.078; EAS: β = 0.32, *P* = 0.0004; SAS: β = 0.14, *P* = 0.013).

### Linked-SFS selection tests

The linked-site frequency spectrum tests (LSFS) is a newly developed family of neutrality tests especially appropriate for the case of inversions, which are a direct application of nearly optimal linear tests for neutrality (Ferretti et al., 2010) to the frequency spectrum of linked sites. We used the simplified version of the tests, i.e. weights were chosen computing the covariance in the approximation of unlinked sites, and we assumed strong selection coefficients in two scenarios: (1) classical selective sweep (positive selection); and (2) long-term balancing selection. The summary statistic used was the frequency spectrum of variants closely linked to the inversion, including information on their linkage pattern (nested or disjoint) with respect to the inverted allele (Ferretti et al., 2018).

To calculate the LSFS, we considered biallelic SNP data from the 1000GP Ph3 in the 434 samples with inversion genotypes. We removed from the analysis all SNPs with a GERP score higher than 2 (Davydov et al., 2010) to reduce the effect of linked selection, as well as SNPs within 0.5 Mb of any of the inversions in our dataset, since their dynamics could be heavily influenced by the inversion itself. We used relatively-small non-overlapping windows of 3 kb in order to reduce the effects of recombination within each window on the empirical null spectrum. The windows tested were localized either within the inversion or the non-recombining flanking regions and skipped the breakpoint interval to avoid errors from associated indels or incorrect short-read mappings. Only 18 autosomal inversions that could be unambiguously phased into the 1000GP haplotypes using perfect tag SNPs within 20 kb from the breakpoints were analyzed. The autosome-wide empirical spectrum was computed on windows of the same size (3 kb) around all autosomal SNPs. Tests were conditioned on the inversion frequency in the different populations. For each test distribution conditioned on minor allele counts of at least 6, a local cubic smoothing was finally applied to the frequency dependence of the distribution, considering derived allele counts between +5 and -5 with respect to that of the inversion. In addition, to control for the complex demographic history of human populations, we used the empirical autosome-wide first and second moments of the empirical linked frequency spectrum of SNPs in each population as a substitute for the null spectrum.

Each population and window was tested separately and population *P*-values of the same windows were combined via Edgington’s method (Edgington, 1972). Combining the results across different windows of an inversion is complicated by the correlation of their *P*-values, since in the absence of recombination they share the same evolutionary history. We dealt with this in two ways. The first approach (conservative) was to assume an arbitrary dependence between windows, and compute the False Discovery Rate (FDR) correcting for multiple correlated testing via Benjamini-Hochberg-Yakutieli (Benjamini and Yekutieli, 2001) for each inversion separately and for all inversions together (in the latter case, HsInv0379 was removed from the analysis due to its size and unbalanced contribution to the statistical noise). The second approach (approximate) is to approximate the joint distribution across correlated windows as a multidimensional Gaussian distribution by: (1) apply a Gaussian transformation to the *P*-values; (2) compute the empirical correlation across all pairs of windows of the same inversion; (3) compute the average Gaussian score for each inversion; (4) build an equicorrelated matrix of the same size as the number of windows in the inversion, with elements equal to 1 on the diagonal and to the empirical correlation off the diagonal; and (5) compare the average Gaussian score with the average score extracted from a multidimensional Gaussian distribution with covariances distributed as the equicorrelated matrix. This approach was applied both to each population separately and to the combined *P*-values from all populations. Inversions with a global conservative *P*-value < 0.01 for either selective scenario (positive or balancing selection) were considered strong selection candidates. Inversions with an approximate *P*-value < 0.05 in at least one population and an approximate or conservative global *P*-value < 0.1 were considered weak selection candidates (Table S8).

### Non-central deviation selection tests

The non-central deviation (NCD) statistics (Bitarello et al., 2018) were adapted to test long-term balancing selection acting on autosomal and chr. X inversion regions. Briefly, NCD1 and NCD2 statistics were computed genome-wide as previously described (Bitarello et al., 2018) in overlapping windows of 2 kb (with 1 kb step) for three target frequencies (0.3, 0.4 and 0.5) using 1000GP Ph3 data from all individuals of the seven populations studied here. A window size of 2 kb was chosen to fit the length of the smaller inversions. After filtering for accessible SNPs according to 1000GP Ph3 pilot accessibility mask, we used as null empirical distribution those windows with a minimum of eight informative sites (either human polymorphic variants in NCD1 and NCD2 or fixed differences with chimpanzee in NCD2 only) and at least 16.7% of positions covered by the hg19-panTro4 alignments available at the UCSC Genome Browser (Kent et al., 2002). Windows of the 44 inversions were defined with the same criteria as in the LSFS test, including the inverted and flanking nonrecombining region, while avoiding breakpoint and IR intervals. Nine inversions did not have any window passing the filtering criteria and were not analyzed (HsInv0031, HsInv0041, HsInv0045, HsInv0055, HsInv0061, HsInv0072, HsInv0344, HsInv0409, and HsInv1124).

A raw empirical *P*-value was assigned to each inversion window corresponding to their quantile in the genome-wide distribution of the statistic computed with the target frequency most similar to the inversion global MAF (Bitarello et al., 2018), and the lowest *P*-value of all the windows for each inversion and population was selected. To correct for the fact that some inversions have more than one window, we then sampled 1,000 sets of regions of equal size and from the same chromosome as each of the inversions, selected the lowest *P*-value of all the windows of each region, and obtained the empirical distribution of minimum-*P*-values equivalent to that of the inversion. Finally, size-corrected *P*-values for each inversion and population were estimated from the quantile in the corresponding minimum-*P*-value distribution. Since balancing selection signals are expected to be shared across multiple populations (Bitarello et al., 2018), we chose as candidates those inversions with three or more populations with size-corrected *P*-values < 0.01 (strong candidates) or *P*-values < 0.05 (weak candidates). The main limitation of the used *P*-values is that, by reducing recombination, inversions may affect the expected empirical distribution. For example, inversions increase variance in the SFS or the age of alleles. Nevertheless, the reduced recombination means stronger effects of background selection, which results in lower levels of diversity and younger alleles, which are the opposite to the signatures detected by the NCD tests. An additional limitation of these tests is that the signatures of balancing selection could be due to any SNP within the windows, rather than the inversion itself. However, the functional effects of the inversion are expected to be much stronger than those of a single nucleotide change.

### Annotation of inversion mutational effects

To determine precisely the direct effect of inversions on genes, we retrieved gene and transcript coordinates from the Comprehensive Gene Annotation Set from GENCODE version 26 (Harrow et al., 2012. We included isoforms from gene annotations with a Transcript Support Level of at least 3, single-exon genes not labelled as “problem”, and pseudogenes. Inversion effects were classified in six different categories (Table S9) according to the following conservative criteria: (1) gene disruption, if there is at least one transcript that encompasses the complete area of one breakpoint; (2) exchange of genic sequences, if two genes of the same family overlap each inversion breakpoint and extend outside of them; (3) inversion of a gene/exon, if the entire gene/exon is situated within the inverted region; (4) inversion of part of an intron, if the inversion and breakpoints are contained inside an intron; and (5) overlap of breakpoints with genes within IRs, if there are genes completely embedded within IRs at the inversion breakpoints. In this last case, due to the high identity of the IRs it is difficult to determine the exact gene effect, although a potential disruption or exchange of gene sequences could happen. If the inversion is associated to indels at the breakpoints, their effect is also taken into account when they include exon sequences. Finally, if none of the criteria above are fulfilled, the inversion is classified as intergenic. We considered inversions affecting directly gene structure (HsInv0102, HsInv0201), disrupting transcripts (HsInv0124, HsInv0340, HsInv0379, HsInv1051) or exchanging gene sequences (HsInv0030) as clear candidates of having functional consequences (Figure 2).

### Gene expression analysis in LCLs

To determine the effect of 42 polymorphic inversions on gene expression in lymphoblastoid cell lines (LCLs), we analyzed 173 individuals experimentally genotyped (42 CEU, 84 TSI and 47 YRI) with data available both in the GEUVADIS project (Lappalainen et al., 2013) and the 1000GP (The 1000 Genomes Project Consortium, 2015). Inversions HsInv0097 and HsInv0379 with MAF <1% in these samples and HsInv0832 located on chr. Y were excluded from the analysis. In addition, to increase the power for discovering gene-expression associations, we imputed our inversions in the complete set of 445 individuals (272 new) from the GEUVADIS Project in common with 1000GP (including 89 CEU, 91 TSI, 86 GBR, 92 FIN and 87 YRI). All these analyses were done with inversion genotype file v.4.7. For 19 unique inversions, unobserved genotypes were inferred through the already identified tag SNPs (*r*^2^ = 1). For the rest, we merged our inversion genotypes and all the variants from the 1000GP Ph3 (including both SNPs and structural variants) for the 173 individuals in common, and used this as reference panel in the imputation. Imputation was performed with IMPUTE v2.3.2 (Howie et al., 2009) adapted to unphased reference genotypes, due to the difficulty of phasing correctly recurrent inversions. We used a region of 1.5 Mb at each side of the inversion and an effective population size of 20,000 (reduced by 25% in chr. X). Genotypes were called with a posterior probability higher than 0.7, and were classified as missing otherwise. We checked the imputation accuracy by removing three random sets of 30 individuals from European (CEU, TSI) and YRI populations from the reference panel, which were subsequently imputed under the same criteria in order to compare imputed and experimental genotypes. Only 14 inversions with an average of correct calls higher than 90% were selected (Figure S4A), resulting in a total of 33 inversions in the imputed dataset.

LCL raw RNA-Seq reads from the GEUVADIS consortium (EMBL-EBI ArrayExpress experiment E-GEUV-1) were aligned against the human reference genome GRCh38.p10 using STAR v2.4.2a (Dobin et al., 2013) and gene-expression levels were estimated as counts and reads per kilobase per million mapped reads (RPKM) based on GENCODE version 26 annotations. Only major chromosomes (chr. 1-22, chr. X, chr. Y and mitochondrial DNA), as well as un-placed and unlocalized scaffolds were considered (patches and alternative haplotypes were not included). Additionally, transcript expression levels were quantified with RSEM v1.2.31 (Li and Dewey, 2011). We only kept the subset of genes and transcripts that has been quantified in at least 20% of the GEUVADIS samples, which results in 29,871 genes and 107,511 transcripts (representing 51% and 54% of the annotated genes and transcripts, respectively). From these, 850 genes and 3,318 transcripts with transcription start sites (TSS) within 1 Mb from inversions were used in the downstream analysis.

*Cis*-acting inversion-eQTL discovery (±1 Mb) was done by two complementary approaches. First, we carried out a targeted study to test only the association between the genotypes of each inversion and gene-expression variation in order to avoid a stringent region-wide multiple-comparison correction. Second, neighboring 1000GP genomic variants and inversions were included in a joint eQTL analysis to estimate the real contribution of each polymorphism to observed gene-expression changes and identify the top variants with highest effect measured by the *P* value (lead eQTLs). *Cis*-eQTLs were mapped through linear regressions implemented in QTLtools v1.1 (Delaneau et al., 2017). First, we ran two separate principal component analysis (PCA) on 1000GP genotype data (considering autosomal variants separated by at least 50 kb with MAF > 0.05) and on RPKM values in order to detect population stratification and capture technical confounding factors that can increase discovery power in association testing. Next, we adjusted expression values in each dataset with a group of covariates comprising individual gender, the first three genotyping principal components of the 1000GP data (corresponding to continent, population and population structure) and technical principal components from the expression PCA (for genes and transcripts, respectively, 10 and 20 in the experimental dataset and 35 and 50 in the imputed dataset). The number of technical covariates was chosen to optimize eQTL finding by maximizing common results and minimizing differences across different eQTL pipelines (see below) but avoiding overfitting the model. This number resulted to be higher for the imputed set, which is concordant with other studies that have shown that it increases with sample size (GTEx Consortium, 2017; Stegle et al., 2012). The individual expression values for each gene were transformed to match normal distributions N(0,1) with QTLtools—normal option (Aulchenko et al., 2007), in order to avoid false positive associations due to any outliers in the data. In the case of the joint eQTL analysis, since there are many variants in *cis* for each gene with different LD patterns, the software allows to account for multiple testing by approximating a null distribution of associations through a permutation approach with a beta distribution. We ran 1,000 permutations to estimate adjusted *P* values for each variant per gene/transcript. eQTL association analysis *P* values were controlled by a significance threshold of 5% false discovery rate (Storey & Tibshirani FDR) (Storey and Tibshirani, 2003) (Table S10) (which was preferred over other multiple comparison correction methods such as Bonferroni that are considered to be overly conservative and give rise to many false negatives (Delaneau et al., 2017)).

We employed different strategies to confirm the reliability of the results. First, we showed that association tests with experimental genotypes and imputed data were highly concordant (Figure S4C). In addition, to check possible biases or inflation of *P* values in the QTLtools analysis, we permuted sample labels from the inversion genotype matrix, maintaining covariates and expression levels. Thus, we broke the potential relationship between inversion genotype and expression, and use this as negative controls. As expected, results from the permuted datasets followed a null distribution of no association (Figure S4D). For inversions located in chr. X, we repeated the analysis excluding heterozygotes to eliminate the effect of the random inactivation of one copy of this chromosome in females. eQTL *P* values were highly correlated in both analysis and just slightly lower when including all samples, with the same significant inversion-gene pairs (Figure S4E), suggesting that the consequences of silencing the chr. X with or without the inversion get averaged across all cells. Finally, we also compared our results with those of two additional commonly-used eQTL mapping methods: the one described by the GTEx Project (GTEx Consortium, 2017) and edgeR-limma (Ritchie et al., 2015; Robinson et al., 2010). In the GTEx analysis, RPKM values were quantile normalized across all samples and gene/transcript expression levels were subsequently adjusted by rank-based inverse normal transformation per each gene and transcript. In this case, technical confounding variation was accounted with the PEER software (Stegle et al., 2012). As mentioned before, to maximize consistent eQTL calls between GTEx and QTLtools pipelines, we tested up to the top 60 expression-derived PEER factors and 60 principal components of the PCA, taken in groups of 5 in decreasing order of the variance explained, and determined the optimal number according to the results overlap. Linear regressions were then done with FastQTL v2.0 (Ongen et al., 2016), including the selected PEER factors (for gene and transcript analysis, respectively, 5 and 20 in the experimental dataset and 35 and 55 in the imputed dataset), gender, and the three population principal components as covariates. In the edgeR-limma workflow, raw read counts were corrected by library size in counts per million. Genes and transcripts that passed the expression-level cutoff (0.1 counts per million in at least two samples) were normalized with trimmed mean of M-values (TMM) (Robinson and Oshlack, 2010) and transformed with voom (Ritchie et al., 2015). Next, limma fit an additive linear model to contrast differentially expressed genes across genotypes, including gender, population and sequencing laboratory as covariates. Other potential batch effects were uncovered with the SVA package (1 and 2 for experimental and imputed sets, respectively) (Leek et al., 2018). All *P* values were corrected by Storey & Tibshirani FDR (Storey and Tibshirani, 2003). As shown in Figure S4F, findings using the different pipelines were highly coincident, although a higher number of significant genes/transcripts were estimated by the GTEx pipeline, suggesting that our chosen method based in PCA and QTLtools is more conservative.

### Inversions as eQTLs in other tissues and conditions

We also explored potential inversion effects on gene expression in other tissues through variants that are in LD with them. In particular, we examined 26 inversions by their highest associated SNP across all populations (*r*^2^ ≥ 0.8) and looked for those in common with SNPs that have been already found to be *cis*-eQTLs in different human tissues by the GTEx project (GTEx Analysis Release v7) (GTEx Consortium, 2017) (Table S11 and Figure S5). Moreover, to find signals of potential effects for additional inversions without highly correlated variants, we extended the analysis to SNPs in moderate LD (*r*^2^ ≥ 0.6) with seven recurrent inversions. To determine if inversions were potentially the causal variants of the gene-expression change, we checked whether those SNPs in highest LD were reported as being the first or second lead eQTL in a specific tissue (Figure S6). Similarly, for recurrent inversions, we checked whether the significance of the expression association was stronger when eQTLs and the inversion were in higher LD.

As an independent replication of these results, we also examined the available gene-expression data from blood samples of ~2000 Estonian individuals obtained by hybridization with Illumina HumanHT-v3.0 Gene Expression BeadChip arrays. In this case, we checked directly the effects of 1541 SNPs that were in high LD (*r*^2^ ≥ 0.8) with 33 inversions either globally (27) or just in Europeans (6). These SNPs were already imputed in Estonian samples based on 1000GP Ph1 variants. In total, six potential inversion-eQTL effects were identified in this study in blood (FDR < 5%): HsInv0006 and *DSTYK;* HsInv0058 and *HLA-E* and *HLA-C;* HsInv0095 and *SPP1;* HsInv0201 and *FBXO38;* and HsInv0209 and *FOLR3*. Of those, five were also found in the GTEx or GEUVADIS data, which represents a good degree of consistency considering the different expression quantification platforms and analysis methods used.

In addition, we applied the same strategy used for GTEx data to study the functional consequences of inversions under other conditions, such as infection. In particular, we linked inversions to immune eQTLs associated to the transcriptional response of primary macrophages to live bacterial pathogens, including Listeria and Salmonella in African-American (AA) (n = 76) and European-American (n = 99) individuals (Nédélec et al., 2016). Of the 26 inversions tested with SNPs in *r*^2^ ≥ 0.8 across all populations, only HsInv0201 showed a clear effect, being in perfect LD in Europeans and Africans with the lead eQTL of gene *SPINK6* expression levels after infection with Salmonella. This result was replicated in a subsequent gene-expression study of human macrophages differentiated from a panel of 123 human induced pluripotent stem cell (iPSC) lines of European origin (Alasoo et al., 2018), where HsInv0201 was completely linked with lead variants for *SPINK6* expression in response to salmonella plus interferon γ plus Salmonella stimulation. The analysis of both datasets also confirmed some of the previous gene-expression effects of HsInv0201 in non-stimulated cells.

### Association of inversions with GWAS SNPs

Another important question is whether polymorphic inversions are linked with SNPs, not just associated to expression changes, but that have been previously connected with specific traits or diseases in human GWAS data. For that, we took advantage of the NHGRI Catalog of published GWAS (release 2017-07-17, v1.0, http://www.ebi.ac.uk/gwas/, which stores a curated collection of the most significant SNP from each independent locus highly associated (*P* < 10^−5^) to a particular disease or phenotype (MacArthur et al., 2017). Nonetheless, many true loci involved in disease susceptibility probably have moderate *P* values. Thus, to obtain a more complete picture of inversions and GWAS signals, hits from the database GWASdb (release 2015-08-19, http://iiwanglab.org/gwasdb, which contains less significant (*P* < 10^−3^) genetic variants (Li et al., 2016), were considered as well. In this case, we filtered by the strongest signal per locus (±100 kb genomic region) to remove redundant entries. As before, to identify which inversions are linked to GWAS hits we crossed these variants with those SNPs in high LD with the inversion (*r*^2^ ≥ 0.8). However, because each study is focused on individuals with different ancestry, we used the specific LD of GWAS signals with inversions in the corresponding population or the closest one available (e.g. TSI for Sardinian, JPT for Japanese, CHB for Han Chinese or Singapore Chinese, and GIH for South

Asian, Indian or Bangladeshi). For individuals from the same population group, we used the LD for the whole continent (e.g. EUR for Ashkenazi, Framingham, British, Caucasian or Hutterite, and EAS for Korean). Finally, if populations from different continents were studied, we used the global LD.

On the other hand, we also investigated if there was enrichment in the number of trait-associated signals in the inversion and flanking regions (±20 kb) compared to what should be expected by chance, that could be attributed to the presence of the inversion. For that, we crossed both GWAS Catalog and GWASdb signals with 1000GP variants, obtaining a final number of 30,720 (from the initial 31,440) and 249,454 (from the initial 251,835) variants in each dataset. Second, GWAS SNPs in high LD (*r*^2^ ≥ 0.8) and associated exactly to the same phenotype were grouped together to obtain a non-redundant set of GWAS signals. To test for a general enrichment of SNP-trait associations surrounding inversions, we carried out a permutation approach, obtaining 1000 random genomic regions as a background model for each inversion. We controlled both for inversion size (defined as the interval comprising the breakpoints ± 20 kb), and SNP frequencies, since control regions followed similar patterns of SNP frequencies (average SNP frequency ± 0.01 and chi-square test *P* > 0.05 for the number of SNPs with MAF <0.2 and ≥0.2 compared to the inversion region). Genome gaps were taken into account and permuted regions could not overlap in order to avoid confounding the analyses. Also, chr. Y was excluded. Because we observed a consistent non-significant (GWAS Catalog) and marginally-significant (GWASdb) enrichment of hits in inversion extended regions, we explored which inversions were driving the signal. Thus, we repeated the same analysis, but using a one-tailed permutation test for each inversion to take into account that many inversion regions present zero GWAS signals.

### Integrative analysis of functional and selection evidence

Overlap of inversion functional and selection evidences was calculated by a Fisher’s exact test of independence. To reduce possible spurious signals, we focused on selection signatures calculated on the inversion itself (excluding NCD1 and NCD2 test results) and all functional evidences except those from GWAS data, which in most cases are related to diseases and could be associated to detrimental effects during evolution. Criteria for classification of strong and weak selection and functional signals are explained the Methods section of each test or in Figure 2. The 21 inversions with perfect tag SNPs that have been included in most analysis comprise all NH inversions, with the exception of HsInv0102, plus HsInv0040.

### Data availability

All data are available in Supplementary Tables and it will be deposited in the InvFEST database (http://invfestdb.uab.cat/) once the paper is published.

### Code availability

Code is available upon request.

---

**Figure S1.**
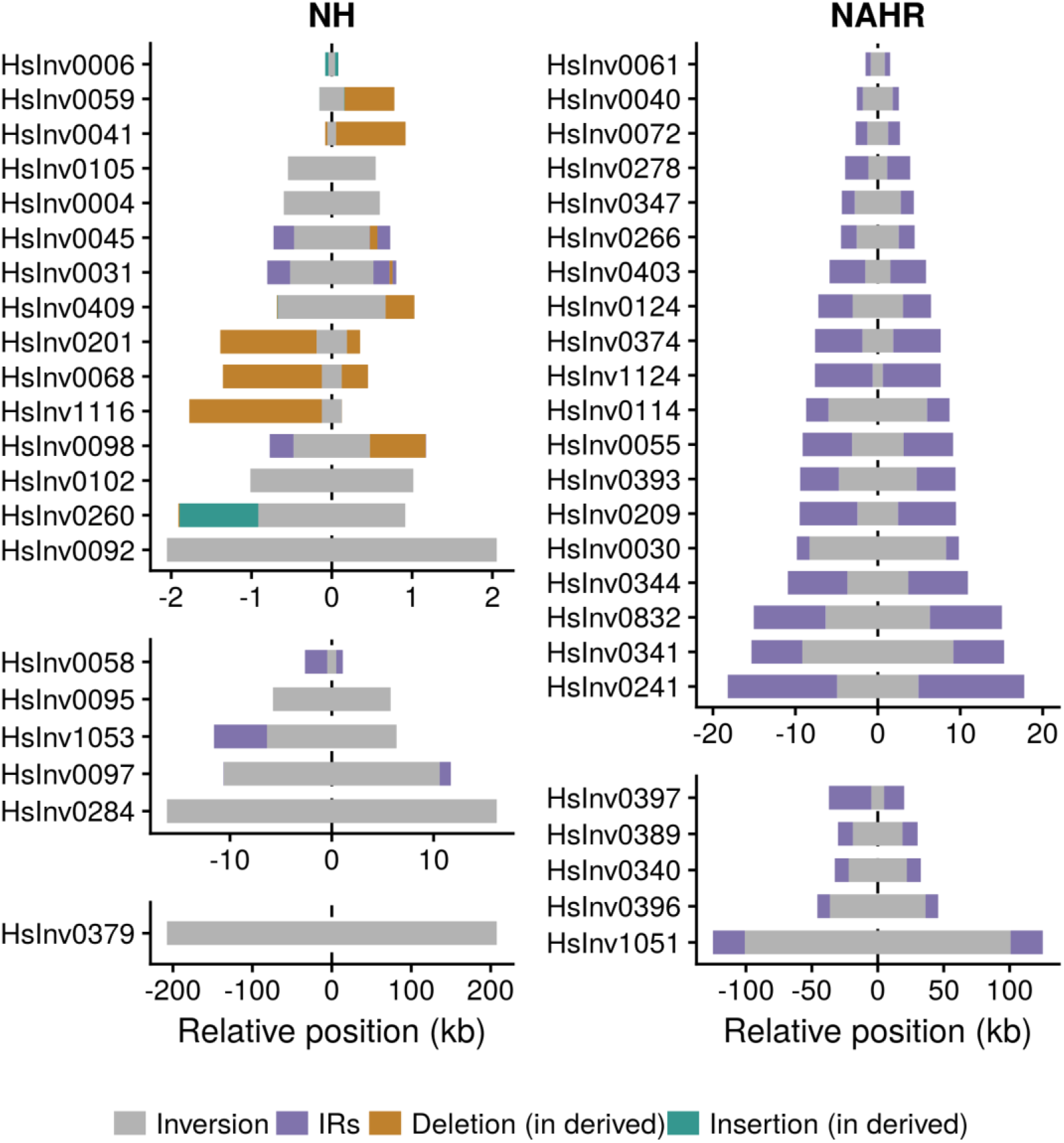
Size and breakpoint complexity of the 45 studied inversions. The graphs illustrate the main characteristics of inversions created by non-homologous mechanisms (NH) or non-allelic homologous recombination (NAHR), with the inverted region represented as a grey bar and flanking inverted repeats (IRs) or other structural changes in different colors. In NH inversions, deletions are sequences present in the original orientation that are eliminated in the derived orientation, and insertions are sequences gained.

**Figure S2.**
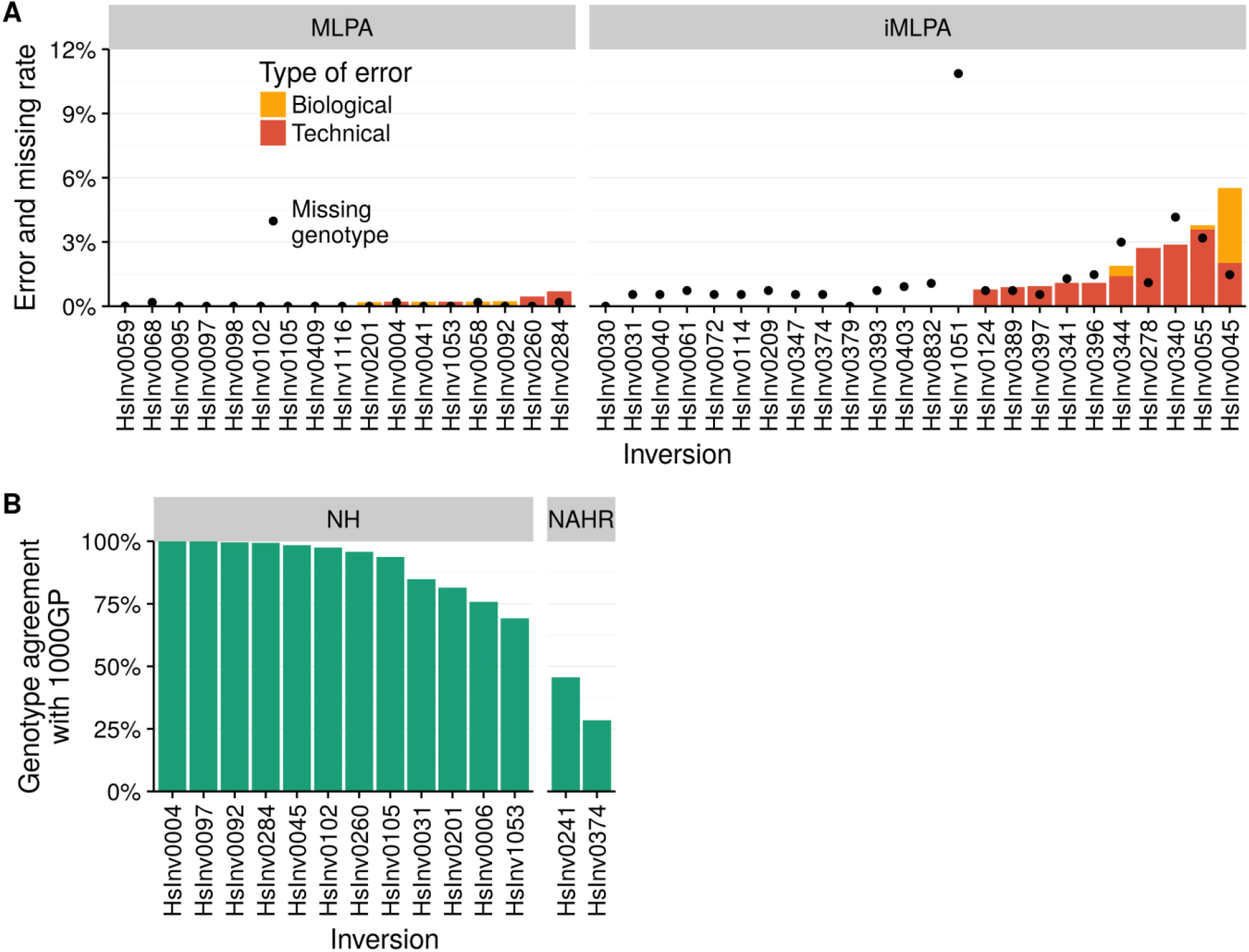
Estimation of inversion genotype accuracy by PCR-based validation and comparison with 1000GP data. A. Genotyping performance of MLPA and iMLPA assays. Inversion genotypes from MLPA/iMLPA were compared with those obtained from PCR or iPCR, plus those imputed from perfect tag SNPs in 1000GP Ph3 data. Genotyping success rate was 99.96% for MLPA and 98.56% for iMLPA. The lower success in iMLPA was due to a lower self-ligation efficiency of large restriction DNA fragments compared to shorter ones (as in the case of HsInv1051), which reduces the amount of specific probe target region and results in smaller amplification peaks, and to problems in specific samples (with one third of missing genotypes accumulating in just three samples). Biological errors correspond to known problems due to restriction site polymorphisms or DNA contamination, whereas technical errors do not have a clear cause. B. Genotype disagreement between the 14 inversions in common with the 1000GP structural variant release (Sudmant et al., 2015) for the 434 samples shared in both datasets. Of the genotypes that differ between studies, in 99.1% the difference is due to 1000GP incorrectly assigning the reference orientation to one of the alleles, whereas according to our experiments it should be the alternative, which leads to underestimating the frequency of the inversion. Also, with a few exceptions, 1000GP error rates tend to be much lower in inversions with clean breakpoints than in those flanked by indels or inverted repeats.

**Figure S3.**
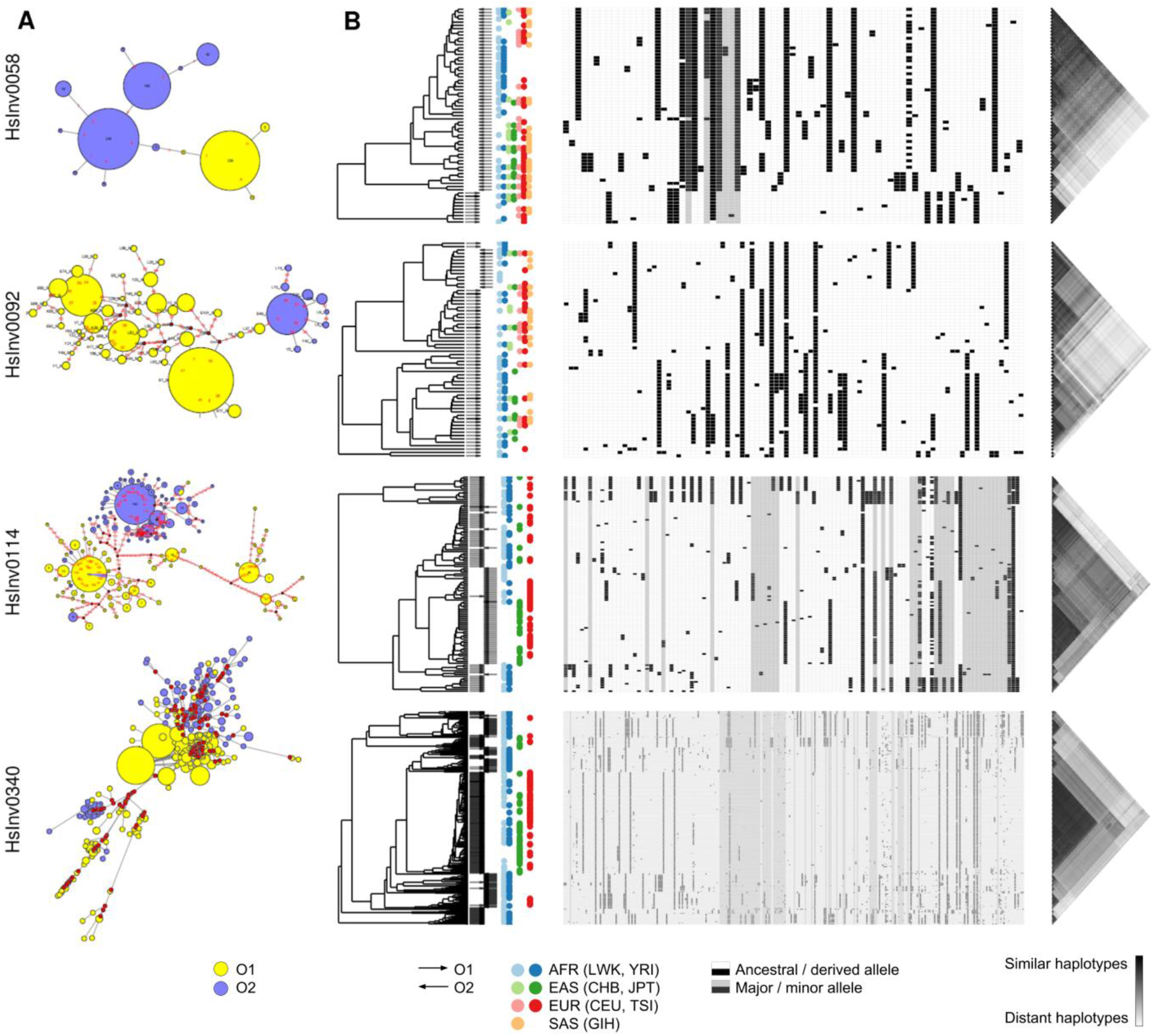
Summary of haplotype relationship for different inversions. A. Representative median-joining networks from 1000GP Ph1 haplotypes obtained with PHASE. Each circle represents a haplotype, whose size is proportional to the number of chromosomes carrying that particular haplotype. Small red points are hypothetical haplotypes not found in the individuals analyzed, and the length of the branch connecting two haplotypes is proportional to the number of changes between them. B. Integrated haplotype plots (iHPlots) for the same four inversions. For unique inversions, the haplotypes correspond to those from 1000GP Ph3 with the extended flanking region whenever possible, whereas for recurrent inversions, the haplotypes are those obtained with PHASE from 1000GP Ph1 data including only the inverted region.

**Figure S4.**
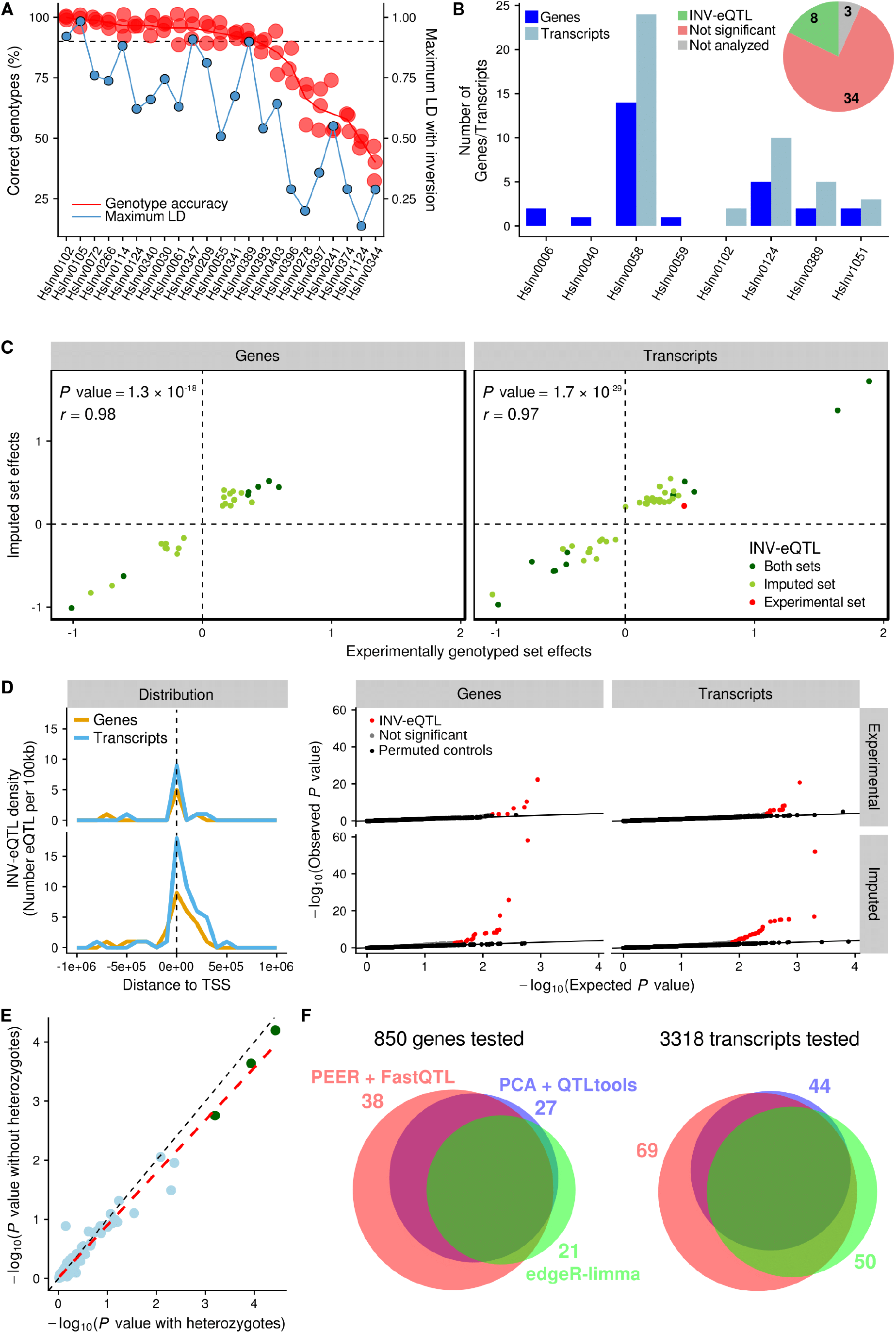
Summary of methodology and main results of inversion gene-expression analysis in lymphoblastoid cell (LCL) data from the GEUVADIS project. A. Genotype imputation accuracy of 23 inversions without perfect tag SNPs based on 1000GP Ph3 variants (SNPs and SVs). The red line represents the mean percentage of right calls in three independent test samples (red dots) of 30 randomly chosen European (CEU and TSI) and YRI individuals with known genotypes that were removed from the reference panel (173 individuals) to validate the imputation. Inversions with at least 90% of masked genotypes correctly imputed (dashed line) were used for gene-expression analysis in other GEUVADIS individuals. Maximum LD between inversions and surrounding genomic variants (blue line) was lower for inversions with worse rates of imputation accuracy. In HsInv0102, which does not have SNPs in high LD, its imputation is based on the 1000GP genotypes for this variant. B. Pie chart summarizing the effects in *cis* of the 45 inversions on LCL expression variation and graph showing the number of genes or transcripts affected by each inversion. Results represented correspond to those from the experimentally-genotyped set (173 individuals) for the 9 inversions that could not be imputed and the extended imputed set (445 individuals) for the 33 imputed inversions. C. Comparison of effect sizes from inversion *cis*-eQTLs (INV-eQTLs) identified in the experimental and imputed sets in LCLs. Concordance of gene and transcript INV-eQTLs between both sets for inversions that could be imputed is shown in different colors: dark green, replicated in both sets; light green, specific of the imputed set; and red, specific of the experimentally-genotyped set. D. *Cis*-eQTL analysis of inversions in LCL expression data. Left: Distribution of INV-eQTLs with respect to the transcription start site (TSS) of the affected genes and transcripts. Inversions tend to locate closer (<100 kb) to genes or transcripts affected compared to all association tests performed both for the experimental (top, *P* = 0.013 and *P* = 0.0005, respectively) and imputed data sets (bottom, *P* = 0.018 and *P* = 3 × 10^−6^, respectively). Right: Quantile-quantile plot of associations between inversions and gene or transcript expression for the experimentally-genotyped and the imputed sets: red dots, significant INV-eQTLs (FDR < 0.05); grey dots, not significant associations; and black dots, negative controls obtained by permuting genotypes relative to covariates and phenotypes, which follow the expected *P* value distribution assuming no-association. E. Correlation of gene eQTL analysis *P* values for inversions located in chr. X with and without heterozygous females. Significant associations (FDR < 0.05) in both analyses are indicated as green dots, with the observed and perfect 1:1 correlation as a red and black dashed line, respectively. F. Results of inversion effects in gene and transcript expression when using different approaches: “PCA+QTLtools”, which corresponds to the pipeline used in this paper (Delaneau et al., 2017) (blue); “PEER+FastQTL”, which corresponds to the pipeline used in the GTEx Project (GTEx Consortium, 2017) (red); and “edgeR-limma” (Ritchie et al., 2015; Robinson et al., 2010) (green). Numbers indicate the significant inversion-gene or inversion-transcript pairs with each analysis method. Venn diagram was done with BioVenn (Hulsen et al., 2008).

**Figure S5.**
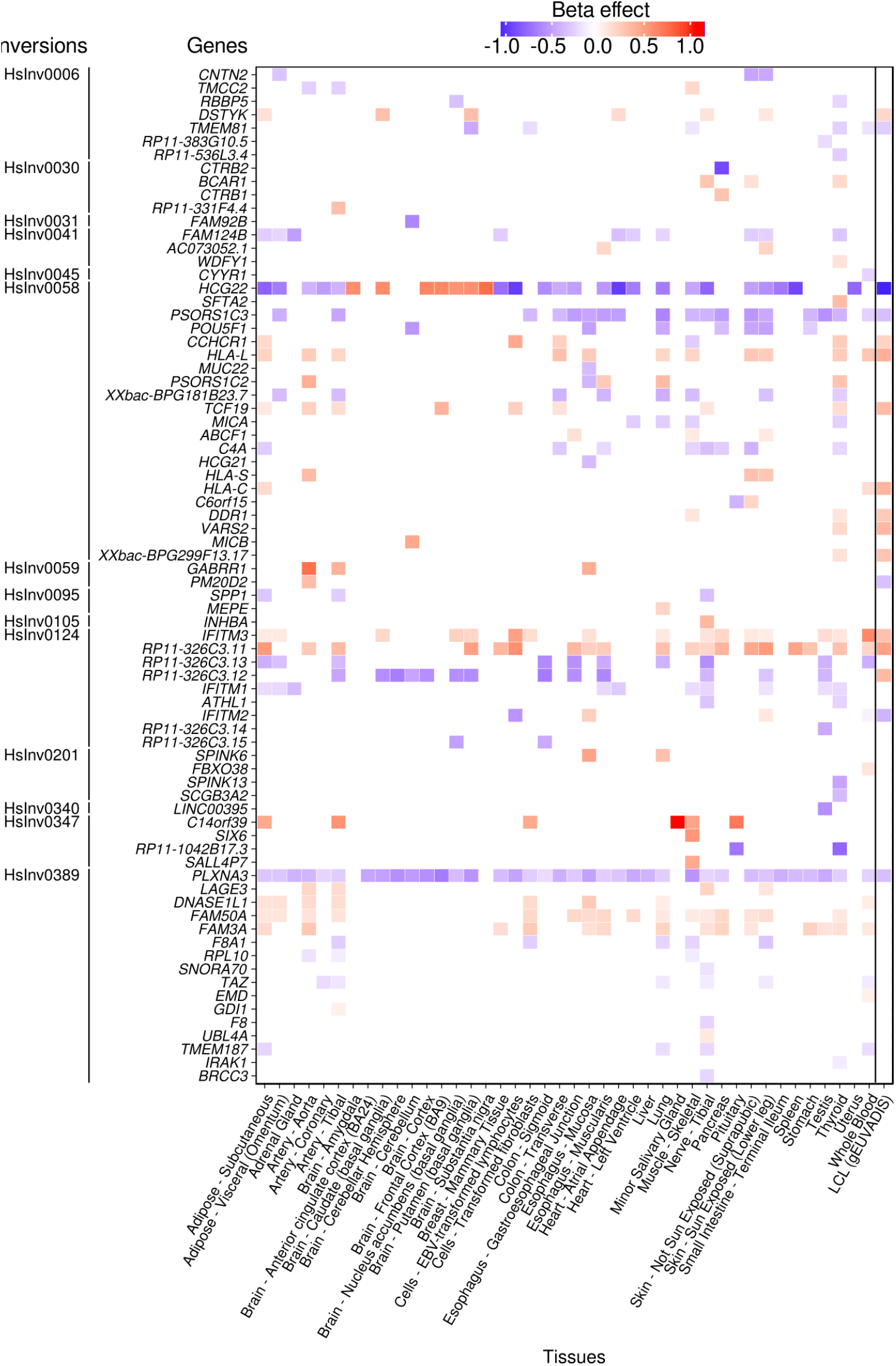
Summary of inversion effects on gene expression across different tissues from the GTEx project. Inversion effects were estimated through variants in high LD (*r*^2^ ≥ 0.8), or moderate LD (*r*^2^ ≥ 0.6) for recurrent inversions, reported as eQTLs in GTEx Analysis Release v7 (see Table S11). The direction and strength of the beta effect of the eQTL is indicated in different color, with blue and red representing respectively lower and higher expression associated to the *O2* orientation of the inversion. Inversion gene eQTLs also identified in the LCL analysis from the GEUVADIS data are represented in the last column.

**Figure S6.**
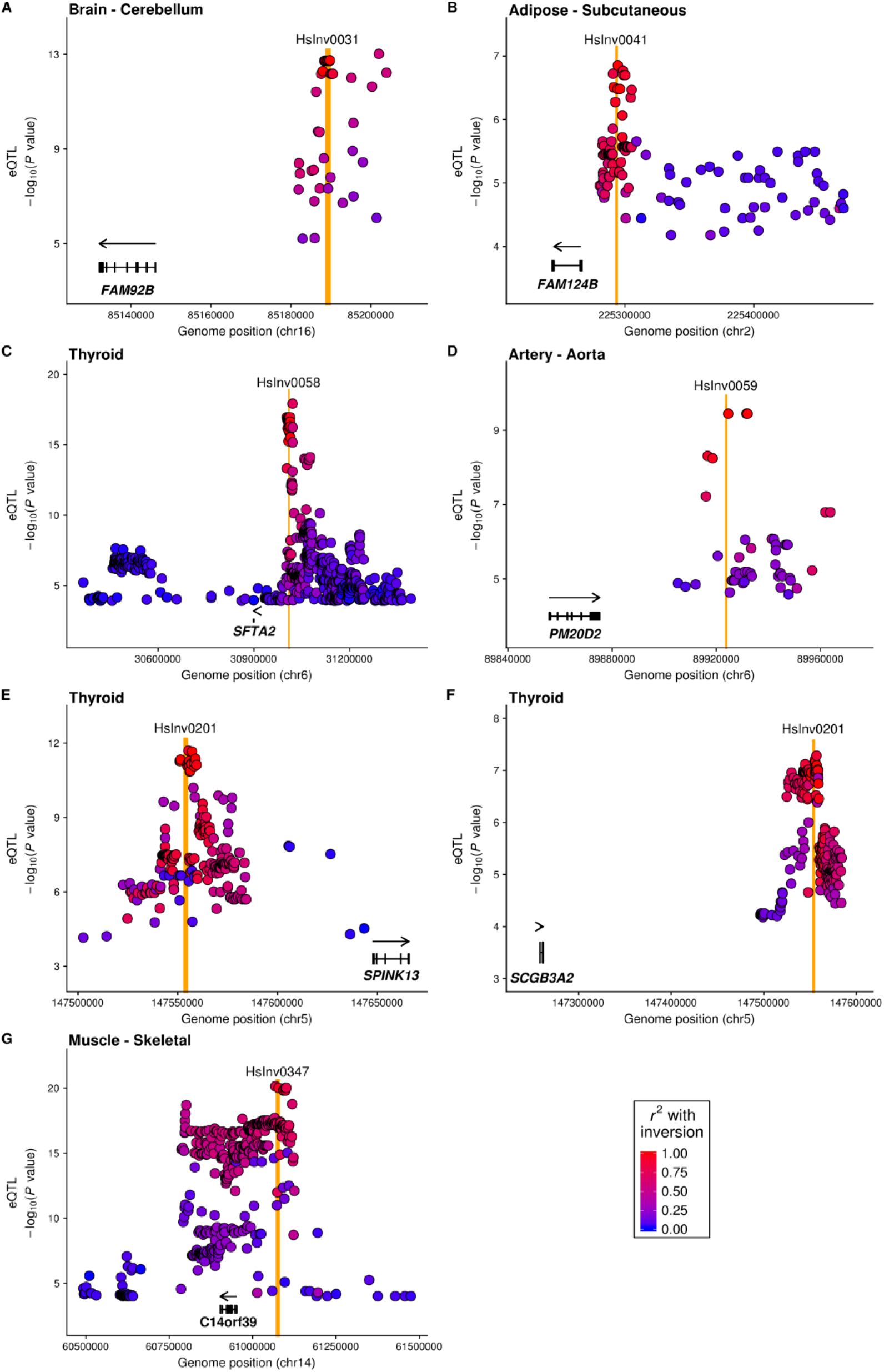
Examples of potential expression effects of inversions in different tissues. Manhattan plots for *cis*-eQTLs associations in tissues from the GTEx project in which an inversion shows the highest LD (*r*^2^ ≥ 0.9) with the two first lead markers. The orange bar pinpoints the inversion position and its LD to each variant is represented in different colors. The affected genes are shown in black and arrowheads indicate the direction of transcription.

**Figure S7.**
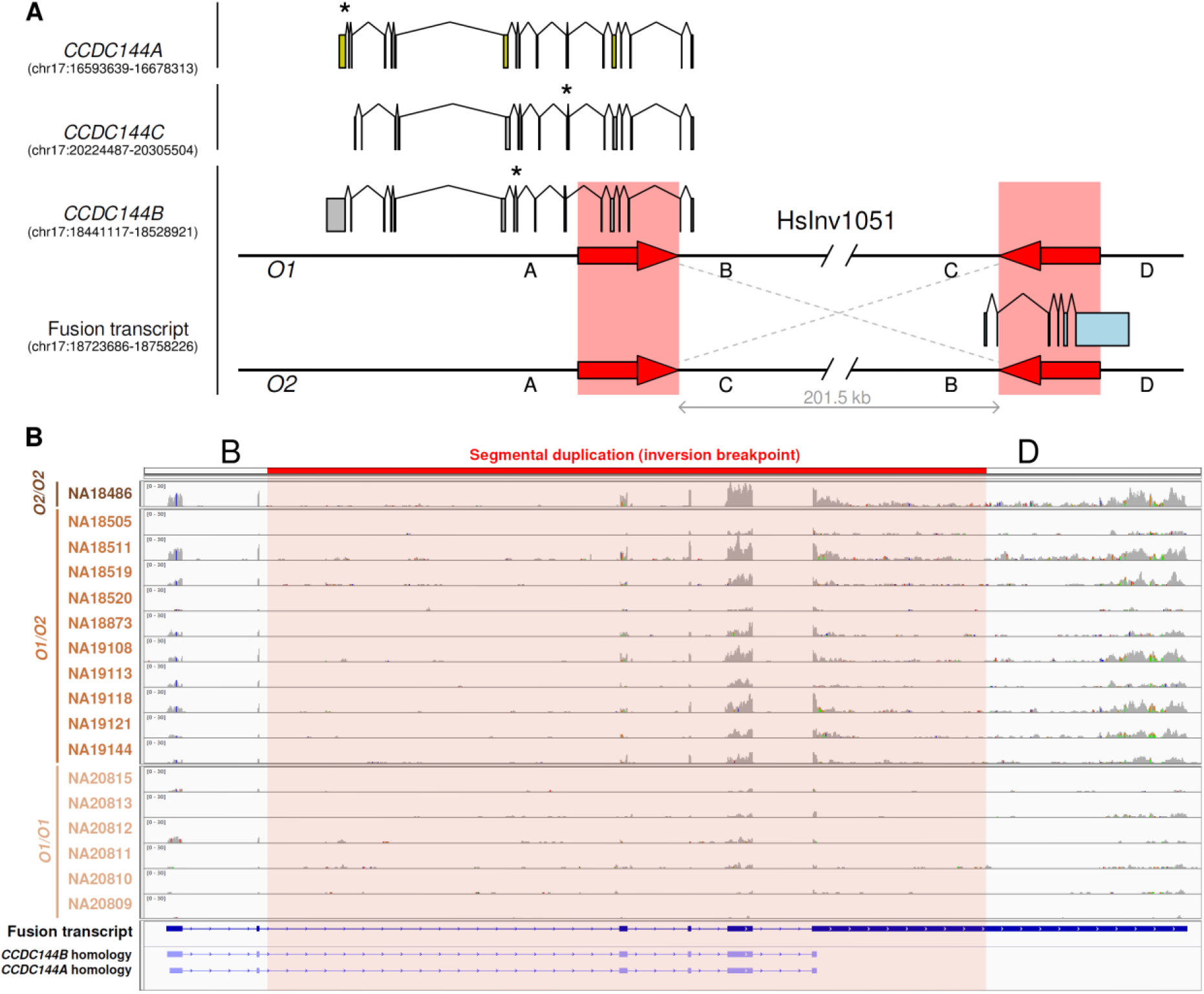
Representation of the fusion transcript created by HsInv1051. A. Diagram of *CCDC144B* gene disruption by inversion HsInv1051 and the novel fusion transcript created including additional 3’ sequences from region D (light blue), with the segmental duplications at the inversion breakpoints represented as red arrows. *CCDC144B* is part of a family with two other members, *CCDC144A* and *CCDC144C*, with 99% identity and very similar exon-intron structure (shown on top). Nevertheless, whereas *CCDC144A* encodes a 1,427-amino acid protein, *CCDC144B* and *CCDC144C* have different frameshift mutations that reduce their coding capacity to 725 and 646 amino acids, respectively (with stop codons shown by asterisks). *CCDC144B* premature stop codon is not be included in the fusion transcript from the inverted allele. B. RNA-Seq profiles from GEUVADIS LCL reads mapped to the inversion BD breakpoint, which was created by reversing *in silico* the sequence between the HsInv1051 breakpoints in the human reference genome (hg19). Reads were remapped to this construct using STAR 2-pass (Dobin et al., 2013) to improve the accuracy of alignments, revealing a novel fusion transcript expressed only in heterozygotes and at higher levels in homozygotes for the inverted allele. The chimeric transcript structure is shown below, after its precise reconstruction with Cufflinks (default parameters) (Roberts et al., 2011) by merging all reads from these samples around the breakpoint region. In addition, its homology with the first six exons of *CCDC144B* and *CCDC144A* is also shown. RNA-seq profiles were visualized on Integrative Genomics Viewer (Robinson et al., 2011).

**Figure S8.**
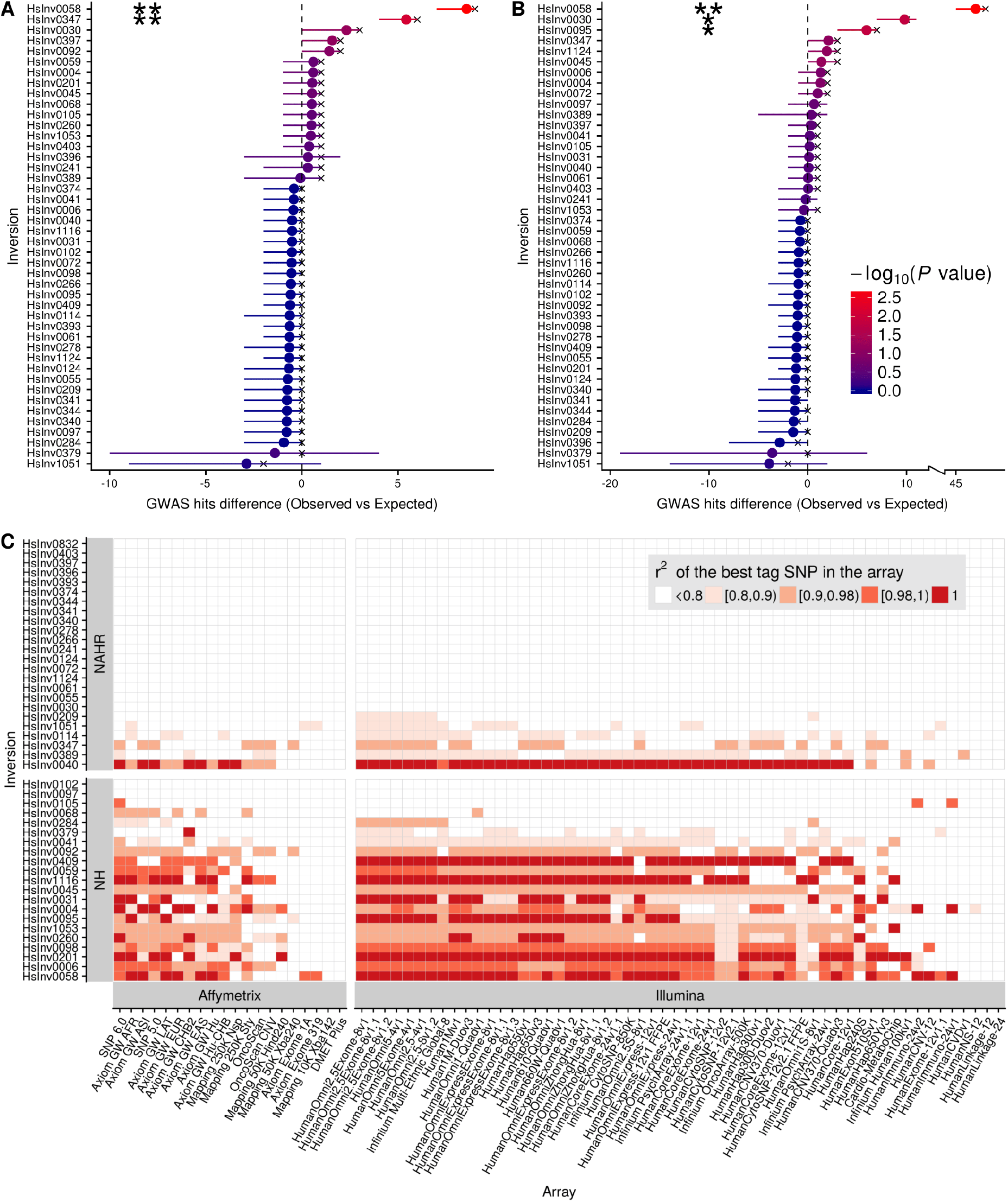
Potential phenotypic effects of inversions from GWAS data. A-B. Enrichment of GWAS signals around 44 autosomal and chr. X inversions (inverted region ± 20 kb) in the GWAS Catalog (A) and GWASdb (B) databases. Error bars show the 0-0.95 interval of the difference in the number of GWAS hits compared with a random background model, together with the mean (filled circle) and the median (cross) of the differences. The color indicates the *P* value of the enrichment according to the scale shown. *, *P* < 0.05; **, *P* < 0.01. C. Coverage of SNPs associated to inversions by checking the presence of inversion global tag SNPs (*r*^2^ > 0.8) in 76 commonly-used genotyping arrays available through the LDLink web portal (Machiela and Chanock, 2015). The LD with the inversion of the best global tag SNP (*r*^2^ > 0.8) in each array is indicated in different colors, showing that for the great majority of NAHR inversions and several of the NH inversions there are not tag SNP or they are not present in the array (represented as white squares). The best performing arrays assessed, HumanOmni5-4v1 and HumanOmni5Exome-4v1 (Illumina), could detect up to 23 inversions (51%), with only 7 being represented by perfect global tag SNPs (*r*^2^ = 1), and 16 by variants with lower LD.

